# The innate immune kinase TBK1 directly increases mTORC2 activity and downstream signaling to Akt

**DOI:** 10.1101/2021.03.30.437707

**Authors:** AS Tooley, D Kazyken, C Bodur, IE Gonzalez, DC Fingar

## Abstract

TBK1 (TANK-binding kinase 1) responds to microbial pathogens to initiate cellular responses critical for host innate immune defense. We found previously that TBK1 phosphorylates mTOR (mechanistic target of rapamycin) (on S2159) to increase mTOR complex 1 (mTORC1) activity and signaling in response to the growth factor EGF and the viral dsRNA mimetic poly(I:C). mTORC1 and the less well studied mTORC2 respond to diverse cues to control cellular metabolism, proliferation, and survival. Here we demonstrate that TBK1 activates mTOR complex 2 (mTORC2) directly to increase Akt phosphorylation at physiological levels of protein expression. We find that TBK1 phosphorylates mTOR S2159 within mTORC2 *in vitro*, phosphorylates mTOR S2159 in cells, and interacts with mTORC2 in cells. By studying MEFs lacking TBK1, as well as MEFs, macrophages, and mice bearing an *Mtor S2159A* knock-in allele (*Mtor^A/A^*), we show that TBK1 and mTOR S2159 phosphorylation increase mTORC2 catalytic activity and promote mTOR-dependent downstream signaling to Akt in response to several growth factors and poly(I:C). While microbial-derived stimuli activate TBK1, we find that growth factors fail to activate TBK1 or increase mTOR S2159 phosphorylation in MEFs. Thus, we propose that basal TBK1 activity cooperates with growth factors in parallel to increase mTORC2 (and mTORC1) signaling. Collectively, these results reveal crosstalk between TBK1 and mTOR complexes (mTORCs), key nodes within two major signaling systems. As TBK1 and mTORCs have each been linked to tumorigenesis and metabolic disorders, these kinases may work together in a direct manner in a variety of physiological and pathological settings.

**One Sentence Summary:** The innate immune kinase TBK1 directly activates mTORC2

## Introduction

TBK1 and its tissue-restricted orthologue IKKε mediate innate immunity against pathogenic viruses and bacteria in response to microbial-derived stimuli (1–5). Viral dsRNA and bacterial LPS bind to and activate the pathogen recognition receptors (PRRs) toll-like receptor 3 (TLR3) and TLR4, respectively (1–5). TLR3/4 activate the kinases TBK1/IKKε through phosphorylation of their activation loop site (S172) by an unknown upstream kinase or through oligomerization and activation loop site auto-phosphorylation (S172) (6–9). TBK1/IKKε in turn phosphorylate the transcription factors IRF3 and 7, resulting in their translocation into the nucleus where they induce expression of type I interferons (e.g., IFNα/β), multi-functional cytokines that initiate host defense responses while limiting tissue damage (10,11).

In prior work, we found that TBK1 phosphorylates mTOR on S2159, which increases mTOR complex 1 (mTORC1) signaling, mTORC1-mediated cell growth (i.e., cell size/mass) and cell cycle progression, and the production of IFNβ (12,13). This positive role for TBK1 in mTORC1 signaling has been confirmed in other studies (14,15). While studying TBK1 and its regulation of mTORC1, we noted that cellular treatment with the TBK1/IKKε inhibitor amlexanox or TBK1 knockout in MEFs reduced phosphorylation of Akt (on S473), an important metabolic kinase and target of PI3K (13). This observation agrees with other studies (14–18), several of which reported that TBK1 phosphorylates Akt S473 directly (16–18). As Akt S473 represents an established target of mTOR complex 2 (mTORC2) (19–22), we investigated more fully the mechanism by which TBK1 promotes Akt phosphorylation, testing the hypothesis that TBK1 directly activates mTORC2 and its downstream signaling to Akt.

The mechanistic target of rapamycin (mTOR) comprises the catalytic kinase core of two known multi-subunit mTOR complexes (mTORCs) (21–23). The scaffolding protein Raptor defines mTORC1 (24,25) while the scaffolding protein Rictor defines mTORC2 (26,27). These mTORCs integrate a diverse array of environmental cues to control cell physiology appropriate for cell type and context. Indeed, aberrant mTORC function contributes to pathologic states including oncogenesis and obesity-linked metabolic disorders (21–23,28). Despite the physiological importance of mTOR, our knowledge of the upstream regulation of mTORCs remain incompletely defined, in particular mTORC2. mTORC1 drives anabolic cellular processes (i.e., protein, lipid, and nucleotide synthesis) in response to the coordinated action of nutrients (e.g., amino acids; glucose), growth factors (e.g., EGF; IGF-1), and hormones (e.g., insulin) to control cell metabolism and promote cell growth (i.e., mass/size) and proliferation (21–23,29). The insulin/IGF-1 pathway represents the best-characterized activator of mTORC1, which utilizes PI3K signaling to Akt, TSC, and Rheb to activate mTORC1 on the surface of lysosomes during nutrient sufficiency (29–31). mTORC1 in turn phosphorylates a diverse set of targets (32,33), with S6K1 T389 phosphorylation serving as a widely employed readout of mTORC1 activity in intact cells.

Growth factors and hormones also activate mTORC2 in a manner that requires PI3K. It is important to note that the upstream regulation of mTORC2 remains significantly less well defined than mTORC1 (21–23,31,34). Recently, the energy sensing kinase AMPK was shown to activate mTORC2 directly to promote cell survival during energetic stress (35). In addition, the stress inducible protein Sestrin2 was shown to activate mTORC2 (36). mTORC2 phosphorylates Akt S473, a widely employed readout of mTORC2 activity in intact cells (19,20,22). Akt functions as a key mediator of PI3K signaling that controls diverse aspects of cell physiology (31,37). Akt activation absolutely requires phosphorylation of its activation loop site (T308) by PDK1. Phosphorylation of its hydrophobic motif site (S473) by mTORC2 activates Akt further to a maximal level and controls substrate preference (38). While not well understood, Akt S473 phosphorylation promotes and/or stabilizes Akt T308 phosphorylation, as increases or decreases in Akt P-S473 often result in correspondingly similar changes in Akt P-T308 (19,35,39). It is important to note that in addition to mTORC2, several other kinases have been identified as Akt S473 kinases including DNA-PK, ATM, and ILK, and more recently TBK1 and IKKε (16–18,40). Functionally, mTORC2 controls cell metabolism, modulates the actin cytoskeleton, and promotes cell survival (21–23,26,27,41).

Beyond its well-known role in innate immunity, TBK1 has been implicated in oncogenesis and metabolic disorders linked to obesity such as type II diabetes, similar to mTOR and Akt (14–17,28,42–54). In oncogenic KRas transformed cells, TBK1 promotes cell proliferation and survival and the growth of tumor explants *in vivo*, with either mTORC1 or Akt suggested as downstream mediators of TBK1 action (14–17,42–46). In addition, elevated expression of the TBK1 orthologue IKKε contributes to breast cancer oncogenesis (44,55). In diet-induced obese mice, adipocyte specific knockout of TBK1 decreases Akt S473 phosphorylation in white adipose tissue, increases insulin resistance and pro-inflammation, and impairs glucose homeostasis (50,54), a phenotype that overlaps with those resulting from adipocyte-specific knockout of Raptor (mTORC1) or Rictor (mTORC2) (28,56–58). Moreover, treatment of obese mice or human patients with amlexanox, or knockout of TBK1, Raptor (mTORC1 partner protein), or S6K1 (mTORC1 substrate) in mouse adipocytes, reduces adiposity and body mass, in part due to increased energy expenditure (50,52,54,59).

To better understand how TBK1 contributes to health and disease, we investigated the molecular mechanism by which TBK1 controls the phosphorylation of Akt. We find that TBK1 phosphorylates mTOR to activate mTORC2 directly, resulting in increased Akt phosphorylation during cellular treatment with growth factors and the innate immune agonist poly(I:C). This work not only elucidates the poorly defined upstream activation of mTORC2, but it improves our understanding of the contribution of TBK1 and mTORCs to physiology and pathologic conditions such as tumorigenesis and obesity-linked metabolic disorders.

## Results

### TBK1 increases mTOR dependent Akt phosphorylation in response to EGF

To elucidate the mechanism by which TBK1 positively controls Akt phosphorylation, we first analyzed TBK1 wild type (TBK1^+/+^) and knockout (TBK1^-/-^) MEFs. TBK1^-/-^ MEFs displayed significantly reduced phosphorylation of Akt S473 (Figure 1A) (AST 478 & graph) across an EGF time course, consistent with prior work (13,16,17). TBK1^-/-^ MEFs also displayed reduced Akt T308 phosphorylation in response to EGF (Figure S1)(CB 92), and the active-site mTOR inhibitor Torin1 ablated both Akt P-S473 and P-T308 (Figures 1A; S1). These data are consistent with mTORC2 functioning as a major Akt S473 kinase (19,20) and with many reports that Akt S473 phosphorylation promotes Akt T308 phosphorylation (19,35,39). Consistent with our prior work (13), TBK^-/-^ MEFs displayed reduced S6K1 T389 phosphorylation, confirming that TBK1 promotes mTORC1 signaling (Figure 1A). To confirm that reduced EGF-stimulated Akt phosphorylation in TBK1^-/-^ MEFs results from loss of TBK1, we stably expressed vector control or Flag-tagged TBK1 in TBK1^-/-^ MEFs by lentiviral transduction followed by puromycin selection. We selected several independent clones in which expression of exogenous Flag-TBK1 matched endogenous TBK1, as our prior work found that overexpression of TBK1 functions in a dominant negative manner to inhibit mTORC1 signaling (13), similar to overexpression of the mTORC1 subunit Raptor, an artifact common for proteins with scaffolding function. Expression of Flag-TBK1 rescued the reduced Akt S473 phosphorylation displayed in TBK1^-/-^ MEFs stimulated with EGF (Figure 1B) (AST 320). Moreover, stable expression of kinase-dead Flag-TBK1 failed to rescue P-Akt S473 (Figure 1C) (AST 93). These results indicate that the kinase activity of TBK1 rather than its scaffolding function promotes Akt phosphorylation. Consistent with this conclusion, the TBK1/IKKε inhibitor amlexanox significantly reduced Akt P-S473 in response to EGF in MEFs (Figure 1D) (AST 450) and HEK293 cells (Figures 1E) (AST 452). Taken together, these results indicate that TBK1 kinase activity promotes mTOR dependent phosphorylation of Akt S473 and T308 during EGF stimulation.

**Figure 1:**
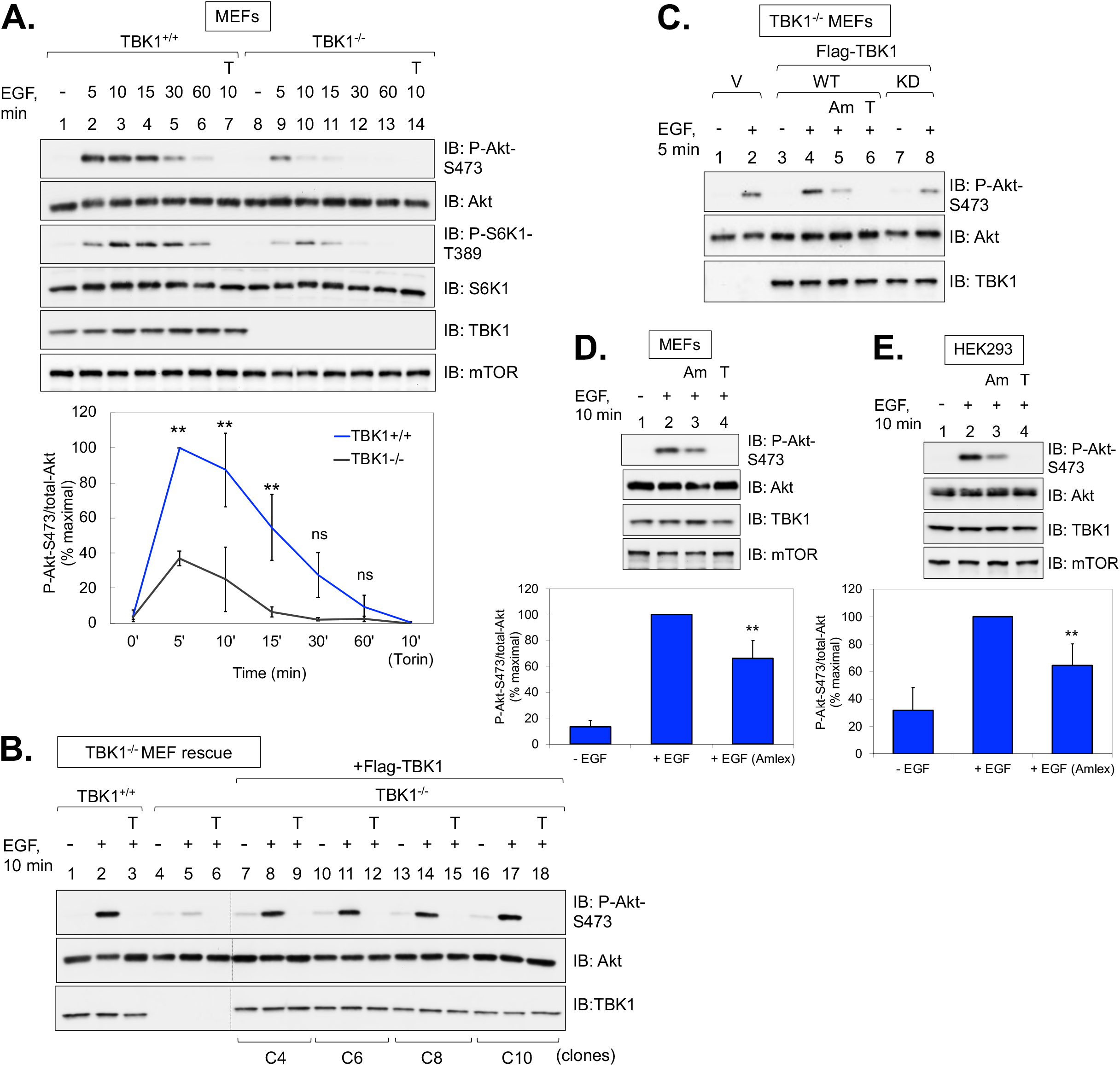
TBK1 promotes mTOR dependent Akt phosphorylation in response to EGF. **A. EGF stimulated Akt S473 phosphorylation in TBK1 null MEFs.** TBK1^+/+^ and TBK1^-/-^ MEFs were serum starved overnight (20 hr), pre-treated with Torin1 (T) [100nM] (30 min), and stimulated without (-) or with EGF [50 ng/mL] for the times indicated times (in minutes, min). Whole cell lysates (WCLs) were immunoblotted with the indicated antibodies. Graph: Quantification of results. Mean ratio +/- SD of Akt P-S473 over total-Akt from three independent experiments, normalized as percent of maximal (+EGF 5 min in TBK1^+/+^ MEFs set to 100%). Statistical significance was measured using paired Student’s t-test (assuming equal variances).**p < .01; “ns”, not significant. **B. Rescue of TBK1 null MEFs with Flag-TBK1.** TBK1^+/+^ MEFs, TBK1^-/-^ MEFs, and clones of TBK1^-/-^ MEFs stably expressing Flag-TBK1 were serum starved, pre-treated with Torin1 [100nM] (30 min), and stimulated without (-) or with (+) EGF [50 ng/mL] for 10 min. WCLs were immunoblotted with the indicated antibodies. **C. Rescue of TBK1 null MEFs with wild type vs. kinase dead Flag-TBK1.** Pools of drug resistant TBK1^-/-^ MEFs stably expressing wild type or kinase dead (K38M) Flag-TBK1 were analyzed as in 1A and B. **D. Effect of amlexanox on EGF stimulated mTORC2 signaling in MEFs.** TBK1^+/+^ MEFs were serum starved overnight (20 hr), pre-treated with amlexanox (Am) [100 μM] (2 hr) or Torin1 (T) [100 nM] (30 min), and stimulated with EGF as in 1A. Whole cell lysates were immunoblotted with the indicated antibodies. Graph: Quantification of results. Mean ratio +/- SD of Akt P-S473 over total-Akt from five independent experiments, normalized as percent of maximal (+EGF 10 min set to 100%). Statistical significance was measured using Student’s paired t-test (assuming equal variances). **p < .01 relative to TBK1^+/+^ MEFs stimulated +EGF in the absence of amlexanox. **E. Effect of amlexanox on EGF stimulated mTORC2 signaling in HEK293 cells.** HEK293 cells were analyzed as in 1E. Graph: Quantification of results. Mean ratio +/- SEM of Akt P-S473 over total-Akt were calculated from five independent experiments as in 1D.

### mTORC2 serves as a major link between TBK1 and Akt S473 phosphorylation at endogenous protein levels

Subsequent to the identification of mTORC2 as the major Akt S473 kinase (19,20,22), several groups demonstrated that TBK1 and IKKε phosphorylate Akt S473 and T308 directly (16–18). As we found that TBK1 phosphorylates mTOR to increase mTORC1 activity and signaling (13), we sought to clarify roles for TBK1 vs. mTOR in phosphorylation of Akt. We therefore assessed effects of mTOR inhibition across an EGF time course. We found that Torin1 ablated Akt S473 phosphorylation and reduced Akt T308 phosphorylation at each time point (1-30 minutes) following EGF stimulation of TBK1^+/+^ and TBK1^-/-^ MEFs (Figure 2A)(DK 324). Consistent with our prior work, TBK1 knockout reduced S6K1 P-T389 (i.e., mTORC1 signaling) and mTOR auto-phosphorylation on S2481 (13). It is important to note that mTOR S2481 auto-phosphorylation represents a simple method to monitor total mTOR or mTORC specific catalytic activity in intact cells (60). These results indicate that in this setting, TBK1 plays a negligible role in Akt S473 phosphorylation, and thus mTOR represents the major Akt S473 kinase in MEFs.

**Figure 2:**
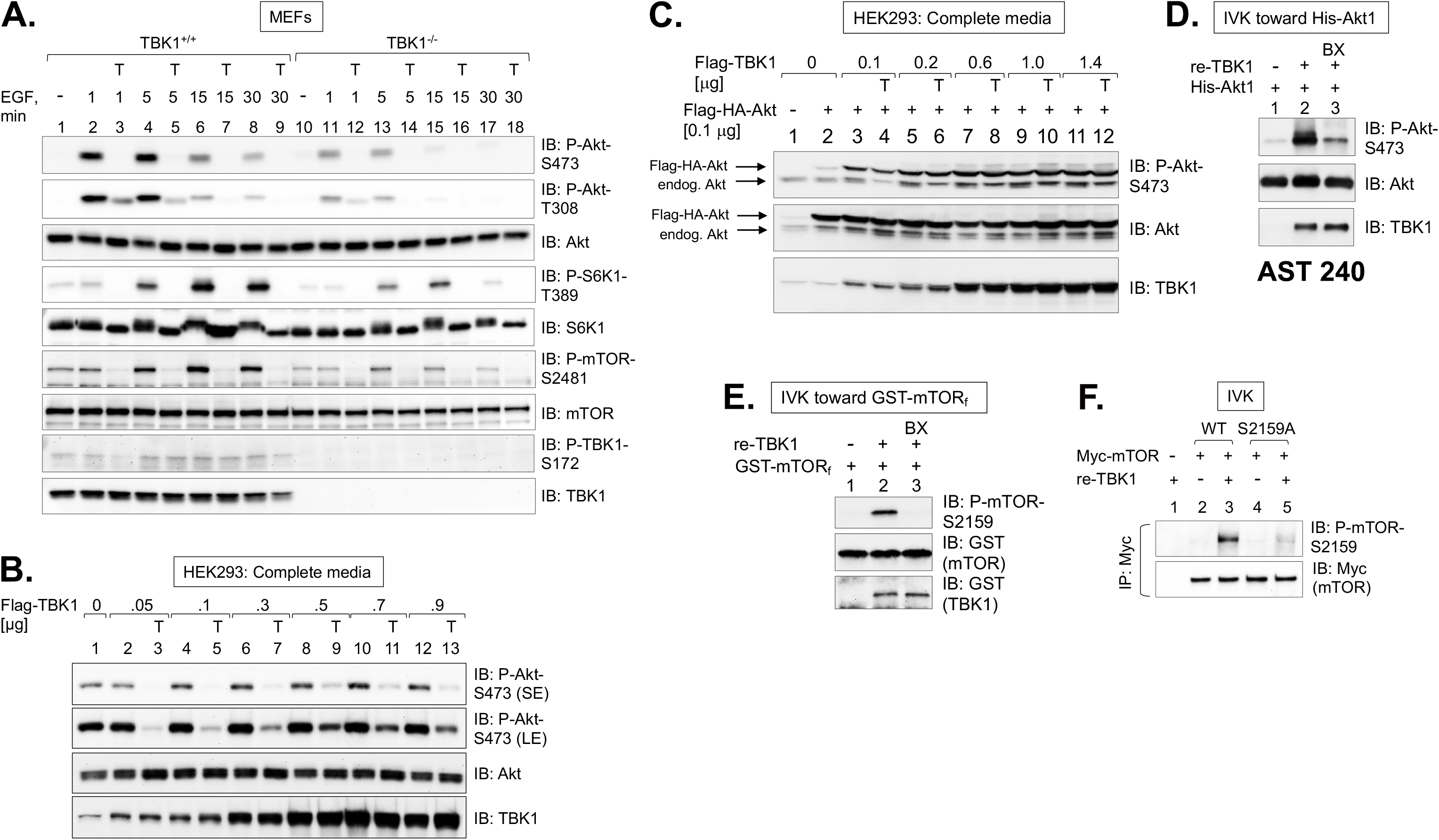
mTORC2 serves as a major link between TBK1 and Akt S473 phosphorylation. **A. Effect of Torin1 on EGF stimulated Akt S473 phosphorylation.** TBK1^+/+^ and TBK1^-/-^ MEFs were serum starved overnight (20 hr), pre-treated with Torin1 (T) [100nM] (30 min), and stimulated without (-) or with EGF [50 ng/mL] for the times indicated (in minutes, min). Whole cell lysates (WCLs) were immunoblotted with the indicated antibodies. SE, short exposure; LE, long exposure. **B. Effect of TBK1 overexpression on Akt P-S473.** HEK293 cells were transfected with increasing amounts of Flag-TBK1 ([0-0.9 μg] per 60 mm plate) in duplicate. ~24 hr post-transfection, cells in complete media were treated with Torin1 (T) [100 nM] (30 min). WCLs were immunoblotted with the indicated antibodies. **C. Effect of TBK1 and Akt co-overexpression on Akt S473 phosphorylation.** HEK293 cells were co-transfected with increasing amounts of Flag-TBK1 ([0-1.4 μg] per 60 mm plate) together with a constant amount of Flag-HA-Akt [0.1 μg] in duplicate. Cells were treated with Torin1 and analyzed as in 1B. **D. Recombinant TBK1 phosphorylates His-Akt1 *in vitro*.** Recombinant, active TBK1 (re-TBK1) [50 ng)]and His-Akt1 [50 ng)]were incubated together in an *in-vitro* kinase (IVK) reaction with ATP at 30°C for 30 min, as indicated. The IVK reaction in lane 3 included pre-treatment with BX-795 (BX) [15 μM] for 30 min prior to initiation of the reaction with ATP. IVK reactions were immunoblotted with the indicated antibodies. **E. Recombinant TBK1 phosphorylates GST-mTORf *in vitro*.** Re-TBK1 [100 ng] was incubated with GST-mTORf [50 ng] at 30°C for 30 min, as indicated. As in 2D, the IVK reaction in lane 3 included pre-treatment with BX-795 (BX) [15 μM]. IVK reactions were immunoblotted with the indicated antibodies. **F. Recombinant TBK1 phosphorylates Myc-mTOR wild type but not S2159A *in vitro*.** HEK293 cells were transfected with vector control (-), Myc-mTOR wild type (WT), or Myc-mTOR S2159A. mTOR was immunoprecipitated (IP) with Myc-9E10 antibody and subjected to IVK reactions with re-TBK1 [50 ng] per reaction, as in 2D and 2E. IVK reactions were immunoblotted with the indicated antibodies.

We next investigated whether elevated levels of TBK1 and/or Akt enables TBK1 to directly engage and phosphorylate Akt. We therefore overexpressed increasing amounts of Flag-TBK1 in HEK293 cells. Consistent with prior work (16–18), exogenous Flag-TBK1 WT increased Akt P-S473 (Figure 2B) (AST 137). Torin1 reduced Akt P-S473 at low to mid doses of Flag-TBK1, indicating mTOR dependency, while higher doses displayed less Torin1 sensitivity. We next co-transfected Flag-TBK1 together with Flag-HA-Akt. The double Flag-HA tag allowed resolution of exogenous Akt (Flag-HA tagged) from endogenous Akt, enabling distinct assessment of phosphorylation on each of these two Akt populations. As before, TBK1 overexpression increased phosphorylation of endogenous Akt in a Torin1 sensitive manner at low doses. Upon co-expression of Flag-TBK1 with Flag-HA-Akt, however, Akt P-S473 became Torin1 resistant at even the lowest dose of Flag-TBK1 (Figure 2C) (AST 239). These data indicate that in the context of TBK1 and Akt over-expression, TBK1 phosphorylates Akt S473 independently of mTOR activity, as reported in prior work (16,17). At physiological levels of protein, however, TBK1 requires mTOR activity to mediate Akt S473 phosphorylation. By *in vitro* kinase assay, we confirmed that TBK1 phosphorylates recombinant Akt1 on S473 (Figure 2D) (AST 240), consistent with other groups (16–18), and TBK1 phosphorylates a recombinant GST-mTOR fragment on S2159 (Figure 2E) (AST 213), consistent with our prior work (13). Importantly, we confirmed that TBK1 phosphorylates wild type but not S2159A Myc-mTOR immunoprecipitated from transfected cells (Figure 2F) (AST 189). Thus, TBK1 phosphorylates diverse substrates with dissimilar consensus phosphorylation motifs, particularly at elevated levels of expression of kinase and/or substrate.

To examine a potential role for TBK1 in the phosphorylation of Akt S473 in the absence of confounding mTORC2 activity, we analyzed MEFs lacking the critical mTORC2 partner protein Rictor. As expected, Rictor^-/-^ MEFs expressing vector control displayed extremely low Akt S473 phosphorylation in response to EGF, while exogenous re-expression of HA-Rictor rescued this phenotype in a manner sensitive to the mTOR inhibitor Ku-0063794 (Figure 3A) (AST 399). Consistent with Figures 1E and 1F, amlexanox inhibited Akt P-S473 in the rescued Rictor^-/-^ MEFs, indicating dependence on TBK1 activity (Figure 3A). With long blot exposure time, however, EGF increased Akt P-S473 in the Rictor^-/-^ MEFs in a Ku-0063794 sensitive but amlexanox resistant manner (Figures 3A, 3B) (AST 399; DK 209). While somewhat unexpected, the Ku-0063794 sensitivity reveals that MEFs lacking Rictor express crippled mTORC2 that still retains a low level of activity toward Akt. In agreement, Xie *et al*. found that mTOR inhibition with Torin1 reduced Akt P-S473 in response to PDGF in Rictor^-/-^ MEFs (17). Therefore, these data indicate that mTOR rather than another kinase (e.g., TBK1) mediates Akt S473 phosphorylation in Rictor^-/-^ MEFs. The amlexanox resistance of Rictor^-/-^ MEFs suggests that TBK1 activity contributes negligibly to Akt phosphorylation in the context of crippled mTORC2. Curiously, shRNA-mediated knockdown of TBK1 in Rictor^-/-^ MEFs reduced Akt P-S473 (Figure 3B) (DK 209), consistent with Xie *et al*. (17). This finding suggests that the scaffolding function of TBK1 may contribute to mTORC2-mediated phosphorylation of Akt S473, at least in cells lacking physiologically intact mTORC2. Taken together, these data indicate that mTORC2 represents a critical link between TBK1 and Akt S473 phosphorylation at physiological levels of protein expression.

**Figure 3:**
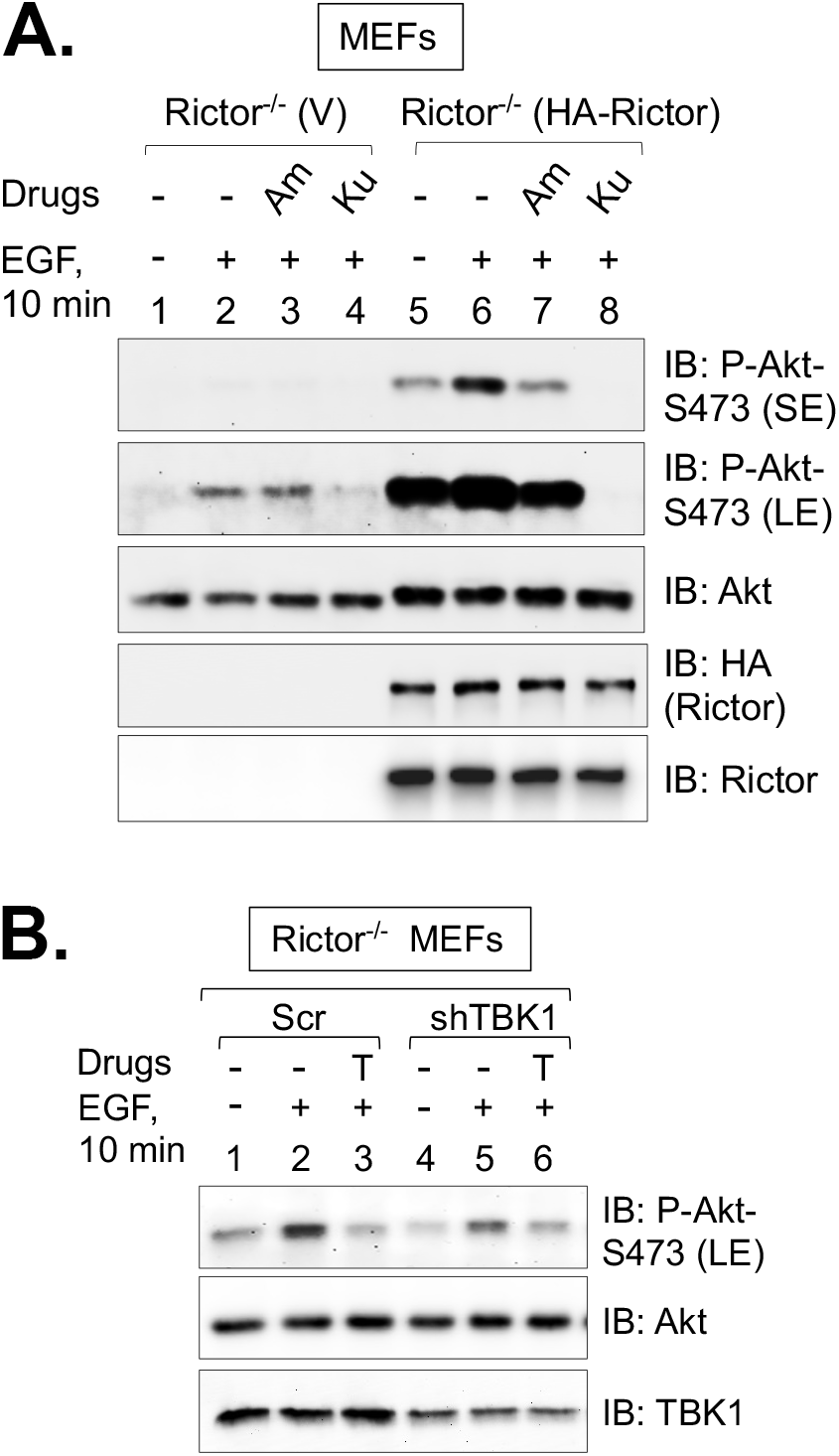
Rictor null MEFs retain a low-level of mTOR dependent Akt S473 phosphorylation supported by TBK1 expression but not TBK1 activity. **A. Effect of Torin1 and amlexanox on Akt S473 phosphorylation in Rictor null MEFs.** Rictor^-/-^ MEFs rescued with vector control (V) or HA-Rictor were serum starved overnight (20 hr), pre-treated with amlexanox (Am) [100 μM] (2 hr) or Ku-0063794 (Ku) [100 nM] (30 min), and stimulated without (-) or with (+) EGF [50 ng/mL] for 10 min. Whole cell lysates (WCLs) were immunoblotted with the indicated antibodies. SE, short exposure; LE, long exposure. **B. Effect of TBK1 knockdown on Akt S473 phosphorylation n Rictor null MEFs.** Rictor^-/-^ MEFs were transduced with lentiviral particles encoding scrambled (Scr) shRNA or an shRNA targeting TBK1 and selected in puromycin. The MEFs were then serum starved overnight (20 hr), pre-treated with Ku-0063794 (Ku) [100nM] (30 min), and stimulated without (-) or with (+) EGF [50 ng/mL] for 10 min. WCLs were immunoblotted with the indicated antibodies.

### mTOR S2159 phosphorylation promotes mTORC2 signaling in response to EGF

In prior work we generated genome edited mice bearing an alanine knock-in substitution at *Mtor S2159* using CRISPR/Cas9 technology (13). By studying bone marrow derived macrophages (BMDMs) in culture isolated from wild type (*Mtor^+/+^*) and S2159A knock-in mice (*Mtor^A/A^*), we demonstrated that mTOR S2159 phosphorylation is required for mTORC1 signaling and IFNβ production in macrophages stimulated with innate immune agonists (i.e., poly(I:C); LPS) (13). Thus, we next investigated a potential direct link between TBK1 and mTORC2 by studying the role of mTOR S2159 phosphorylation in control of mTORC2 signaling. To do so, we isolated littermate-matched MEFs from *Mtor^+/+^* and *Mtor^A/A^* mice (pair #1 MEFs), subjected them to spontaneous immortalization, and analyzed their response to EGF following serum deprivation. Relative to MEFs from *Mtor^+/+^* mice (i.e., mTOR^+/+^ MEFs), MEFs from *Mtor^A/A^* mice (i.e., mTOR^A/A^ MEFs) displayed significantly reduced Akt P-S473 (Figure 4A) (AST 467; graph) and Akt P-T308 (Figure S2A) (AST 466) across an EGF time course. mTOR^A/A^ MEFs also displayed reduced S6K1 P-T389 and mTOR S2481 autophosphorylation (Figure 4A). Activation of the MAPK/ERK pathway in response to EGF remained unperturbed in the mTOR^A/A^ MEFs (as monitored by the phosphorylation of ERK1 and 2 on T202/Y204), indicating intact activation of EGF-receptor signaling to MAPK/ERK in mTOR^A/A^ MEFs (Figure 4A). Importantly, we observed reduced Akt P-S473 and S6K1 P-T389 in response to EGF in a second, independently derived pair of mTOR^+/+^ and mTOR^A/A^ MEFs (pair #2 MEFs)(Figure 4B)(AST 254). These data demonstrate that mTOR S2159 phosphorylation increases mTOR catalytic activity and promotes mTORC2 and 1 signaling upon cellular stimulation with EGF.

**Figure 4:**
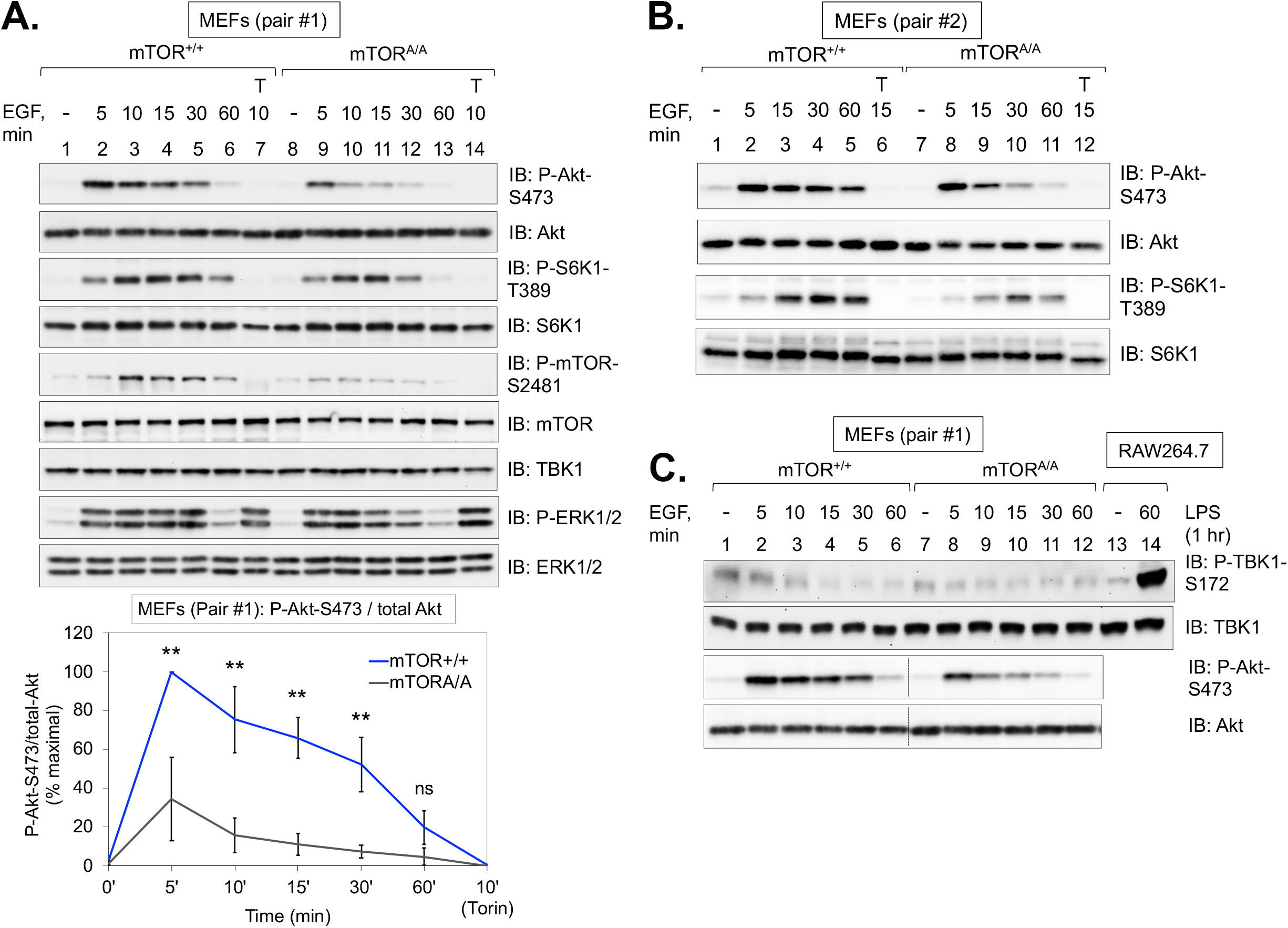
mTOR S2159 phosphorylation promotes mTORC2 signaling in response to EGF. **A. Effect of EGF on Akt P-S473 in mTOR S2159A knock-in MEFs.** Immortalized mTOR^+/+^ and mTOR^A/A^ MEFs (pair #1) were serum starved overnight (20 hr.), pre-treated with Torin1 (T) [100nM] (30 min.), and stimulated without (-) or with EGF [50 ng/mL] for the times indicated (in minutes, min). Whole cell lysates (WCLs) were immunoblotted with the indicated antibodies. Graph: Quantification of results. Mean ratio +/- SD of Akt P-S473 over total-Akt from four independent experiments, normalized as percent of maximal (+EGF 5 min. in mTOR^+/+^ MEFs set to 100%). Statistical significance of differences was measured using Student’s paired t-test (assuming equal variances). **p < .01; “ns”, not significant. **B. Effect of EGF on Akt P-S473 in a second immortalized pair of wild type and mTOR S2159A knock-in MEFs.** Immortalized mTOR^+/+^ and mTOR^A/A^ MEFs (pair #2) were treated as in 4A. **C. Effect of EGF on phosphorylation of the TBK1 activation loop site (S172) in mTOR MEFs.** mTOR^+/+^ and mTOR^A/A^ MEFs (pair #1) were serum starved and stimulated with EGF for the times indicated, as in 4A. RAW264.7 macrophages in complete media were stimulated without (-) or with (+) LPS [100 ng/mL] (60 min.) to serve as a positive control for TBK1 P-S172 western blotting. WCLs from mTOR MEFs and RAW264.7 macrophages were resolved on the same gel and immunoblotted with the indicated antibodies.

We next asked whether EGF activates TBK1 by monitoring TBK1 phosphorylation on its activation loop site (S172). We found that EGF failed to increase TBK1 P-S172 in either mTOR MEFs (Figure 4C) (AST 467-428) or TBK1 MEFs; as expected, LPS treatment of RAW264.7 macrophages strongly increased TBK1 P-S172 (Figure 4C) (Figure S2B) (AST 360-428). Note that EGF also failed to increase P-TBK1 in HEK293 cells (see (13)). These results demonstrate that EGF does not activate TBK1, at least in MEFs and HEK293 cells, and thus basal rather than EGF stimulated TBK1 activity supports mTORC1/2 signaling.

### TBK1 phosphorylates mTOR within mTORC2, interacts with mTORC2, and increases mTORC2 intrinsic catalytic activity

To further define the mechanism by which TBK1 promotes mTORC2 signaling, we asked whether recombinant TBK1 (re-TBK1) phosphorylates mTOR S2159 within mTORC2 directly. It is important to note that our prior work demonstrated that TBK1 phosphorylates mTOR S2159 within mTORC1 (13). By *in vitro* kinase assay, we found that re-TBK1 increased mTOR P-S2159 on Rictor-associated mTOR immunoprecipitated from cells, and inclusion of the TBK1/IKKε inhibitor BX-795 *in vitro* blocked this increase (Figure 5A) (AST 251). We next asked whether TBK1 and mTORC2 interact in cells. By co-immunoprecipitating endogenous proteins, we found that Rictor immunoprecipitates pulled down TBK1 in TBK1^+/+^ but not TBK1^-/-^ MEFs (Figure 5B) (AST 492). Together, these data support a mechanism whereby TBK1 interacts with mTORC2 and subsequently mediates the direct phosphorylation of mTOR S2159 within mTORC2. We next asked whether EGF increases mTOR S2159 phosphorylation and whether TBK1 promotes mTOR S2159 phosphorylation in intact cells. EGF failed to increase P-S2159 on mTOR immunoprecipitated from wild type TBK1^+/+^ or mTOR^+/+^ MEFs (Figures 5C, 5D) (DK 957; DK 959). Importantly, TBK1 knockout from MEFs reduced mTOR P-S2159 (Figure 5C) (DK 959), and mTOR P-S2159 was undetectable in mTOR^A/A^ MEFs (Figure 5D) (DK 957), confirming *Mtor S2159A* knock-in. We speculate that the remaining mTOR P-S2159 found in TBK1^-/-^ MEFs results from IKKε-mediated phosphorylation of mTOR. Indeed, while IKKε expression is generally tissue-restricted and extremely low in non-immune cells, MEFs indeed express detectable levels of IKKε (Figures 5C, 5D). These data demonstrate that TBK1 mediates mTOR S2159 phosphorylation *in vitro* and in intact cells and support the conclusion that basal rather than EGF stimulated TBK1 kinase activity mediates mTOR P-S2159 to promote mTORC2 signaling.

**Figure 5:**
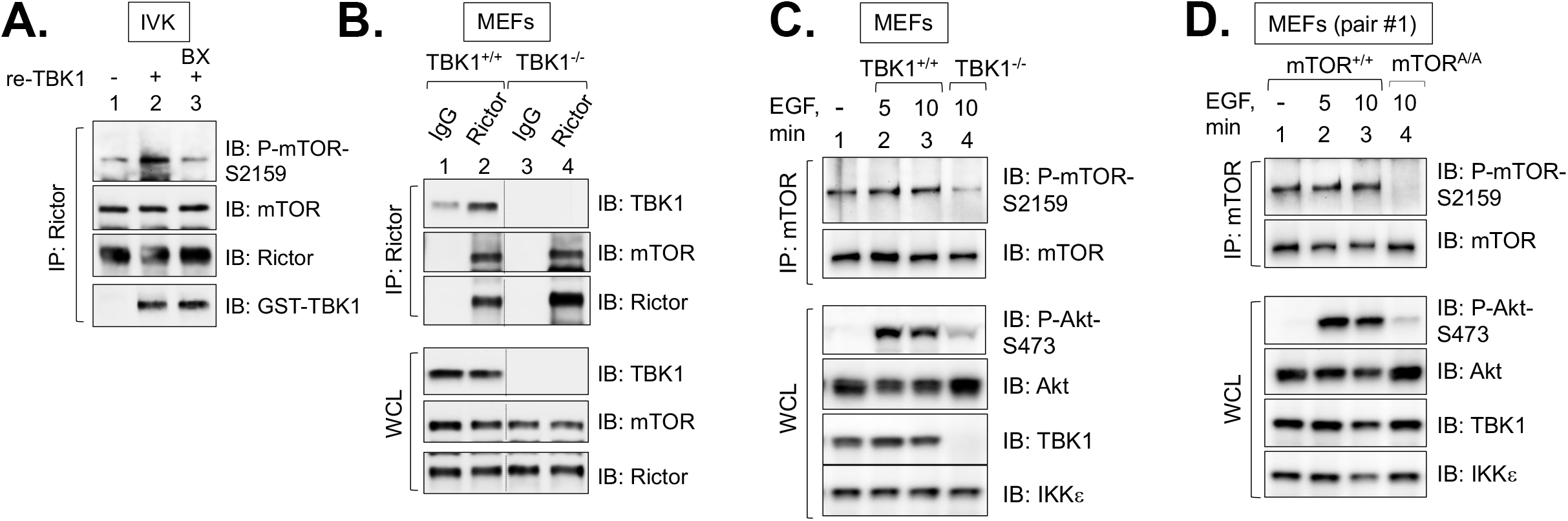
TBK1 phosphorylates mTOR and interacts with mTORC2. **A. Phosphorylation of mTOR S2159 within mTORC2 by TBK1 *in vitro*.** Rictor was immunoprecipitated from HEK293 cells and incubated with recombinant, active TBK1 (re-TBK1) [100 ng] for 30 min.at 30°C. The IVK reaction in lane 3 was pre-treated with BX-795 (BX) [15 μM] for 30 min as indicated. **B. Co-immunoprecipitation of TBK1 with mTORC2 (Rictor/mTOR).** Whole cell lysates from TBK1^+/+^ and TBK1^-/-^ MEFs cultured in complete media (DMEM/FBS) were incubated with Sepharose beads conjugated to either control IgG or anti-Rictor antibodies overnight at 4°C. The immunoprecipitates (IPs) and whole cell lysates (WCLs) were immunoblotted with the indicated antibodies. **C. mTOR S2159 phosphorylation in TBK1 null MEFs.** mTOR was immunoprecipitated from TBK1^+/+^ and TBK1^-/-^ MEFs that had been serum starved overnight and stimulated with EGF [25 ng/mL] (10 min). IPs and WCLs were immunoblotted with the indicated antibodies. **D. mTOR S2159 phosphorylation in mTOR S2159A knock-in MEFs.** mTOR was immunoprecipitated from mTOR^+/+^ and mTOR^A/A^ MEFs as in 5C. IPs and WCLs immunoblotted with the indicated antibodies.

We next asked whether TBK1 and mTOR S2159 phosphorylation increase the intrinsic catalytic activity of mTORC2 by *in vitro* kinase assay. To do so, we immunoprecipitated Rictor from TBK1^+/+^ vs. TBK1^-/-^ MEFs and from mTOR^+/+^ vs. mTORAA MEFs after EGF stimulation of serum deprived cells. The Rictor immunoprecipitates were washed, incubated in kinase buffer with ATP and recombinant His-Akt1 as substrate, and the ability of Rictor-associated mTOR to phosphorylate Akt S473 *in vitro* was monitored by western blot. In both TBK1^+/+^ MEFs (Figure 6A) (DK 894) and mTOR^+/+^ MEFs (Figure 6B) (AST 521), EGF increased mTORC2 catalytic activity in a Torin1-sensitive manner, as expected. The fold increase in mTORC2 catalytic activity mediated by EGF, however, was reduced in TBK1^-/-^ MEFs (Figure 6A) and mTOR^A/A^ MEFs (Figure 6B). To assess the role of TBK1 and mTOR S2159 phosphorylation in control of mTORC2 catalytic activity by an independent approach, we monitored S2481 auto-phosphorylation on Rictor associated mTOR (i.e., mTORC2). Similar to results obtained with mTORC2 *in vitro* kinase assays, the fold increase in mTOR S2481 autophosphorylation mediated by EGF was reduced in TBK1^-/-^ (Figure 6C) (CB 120) and mTOR^A/A^ MEFs (Figure 6D) (AST 506). Taken together, these results indicate that TBK1 and mTOR S2159 phosphorylation increase mTORC2 catalytic activity.

**Figure 6:**
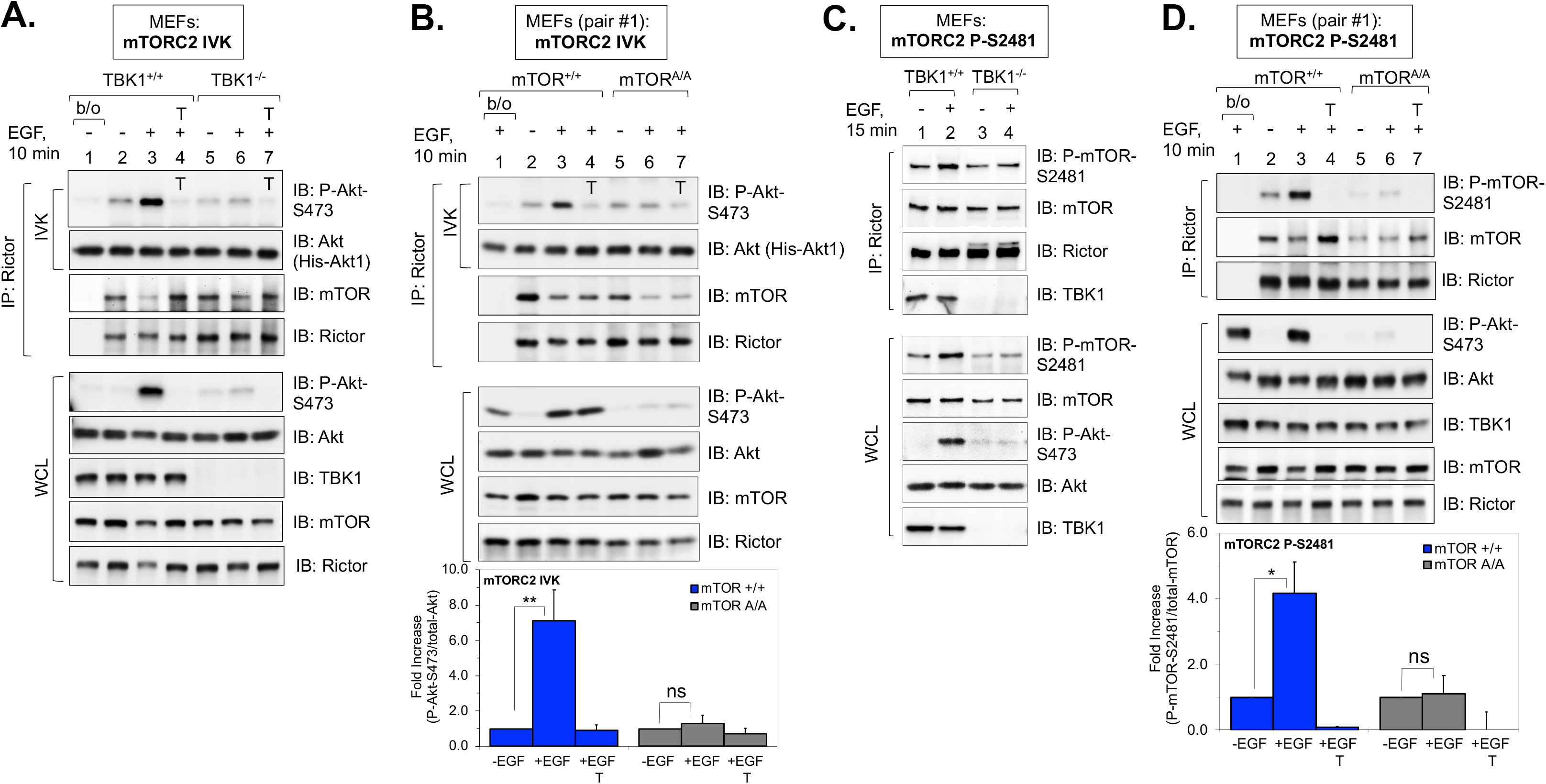
TBK1 and mTOR S2159 phosphorylation increase mTORC2 catalytic activity. **A. Effect of EGF on mTORC2 intrinsic catalytic activity in TBK1 null MEFs.** Rictor was immunoprecipitated (IP) from TBK1^+/+^ and TBK1^-/-^ MEFs that had been serum starved overnight, pretreated with Torin1 [100 nM] (30 min), and stimulated with EGF [50 ng/mL] (10 min). The immune complexes were washed and subjected to *in vitro* kinase (IVK) reactions with His-Akt1 [100 ng] substrate and ATP [250 μM] at 30°C for 30 min. Torin1 was included in the IVK reactions from cells pre-treated with Torin1, as indicated (T on the blot). IVKs and whole cell lysates (WCLs) were immunoblotted with the indicated antibodies. **B. Effect of EGF on mTORC2 intrinsic catalytic activity in mTOR S2159A knock-in MEFs.** Rictor was immunoprecipitated from mTOR^+/+^ and mTOR^A/A^ MEFs (pair #1) that had been treated with EGF. IVK reactions were performed on the immune complexes and analyzed as in 6A, except that only certain IVK reactions but not the cells were treated with Torin1, as indicated (T on the blot). Graph: Quantification of results. Mean ratio +/- SEM of the fold increase in Akt P-S473 over total-Akt1 from four independent experiments, normalized within each genotype, setting the − EGF condition to 1.0. Statistical significance was measured using paired Student’s t-test (assuming equal variances). **p < .01 relative to mTOR^+/+^ MEFs stimulated +EGF; “ns”, not significant. **C. Effect of EGF on mTORC2-specific mTOR S2481 autophosphorylation in TBK1 null MEFs.** Rictor was immunoprecipitated from TBK1^+/+^ and TBK1^-/-^ MEFs that had been serum starved overnight and stimulated with EGF [50 ng/mL] (15 min) as in 6A. IPs and WCLs were immunoblotted with the indicated antibodies. **D. Effect of EGF on mTORC2-specific mTOR S2481 autophosphorylation in mTOR S2159A knock-in MEFs.** Rictor was immunoprecipitated from mTOR^+/+^ and mTOR^A/A^ MEFs (pair #1) that had been treated with EGF and Torin1 as in 6A. IPs and WCLs were immunoblotted with the indicated antibodies. Graph: Quantification of results. Mean ratio +/- SEM of the fold increase in mTOR P-S2481 over total-mTOR from three independent experiments. *p < .05 relative to mTOR^+/+^ MEFs stimulated +EGF; “ns”, not significant.

### TBK1 and mTOR S2159 phosphorylation increase mTORC2 (and mTORC1) signaling in response to diverse growth factors

We next investigated whether the positive role of mTOR S2159 phosphorylation in mTORC2 signaling extends to a broader set of growth factors. We thus interrogated mTORC2 signaling to Akt, as well as mTORC1 signaling to S6K1, in mTOR^+/+^ vs. mTOR^A/A^ MEFs in response to several growth factors known to activate mTORC2 and 1 signaling. We first assessed the role of mTOR P-S2159 in control of mTORC2 and 1 signaling in MEFs cultured in complete media (i.e., DMEM/FBS) containing serum growth factors. mTOR^A/A^ MEFs as well as TBK1^-/-^ MEFs displayed reduced Akt S473 and S6K1 T389 phosphorylation (Figure 7A) (DK 964; AST 287). Treatment of cells with Torin1 ablated these phosphorylation events, indicating that mTOR activity is required for mTORC2 and 1 signaling in complete media (Figure 7A). We next assessed the role of mTOR P-S2159 in control of mTORC2 and 1 signaling in response to stimulation of MEFs with fetal bovine serum (FBS), platelet-derived growth factor (PDGF) (a major constituent of FBS), and insulin (which acts through IGF-1 receptors in MEFs) following serum deprivation. Similar to their response to EGF, mTOR^A/A^ MEFs displayed reduced Akt P-S473 and S6K1 P-T389 in response to all three growth factors with varying dynamics across a time course (Figures 7B-D) (AST 575; AST 574; CB 329). In addition, all three growth factors failed to increase TBK1 S172 phosphorylation (Figure S3), suggesting that growth factor receptor signaling does not activate TBK1, at least in MEFs. These results indicate that mTOR S2159 phosphorylation increases mTORC2 and 1 activity in parallel to growth factor signaling.

**Figure 7:**
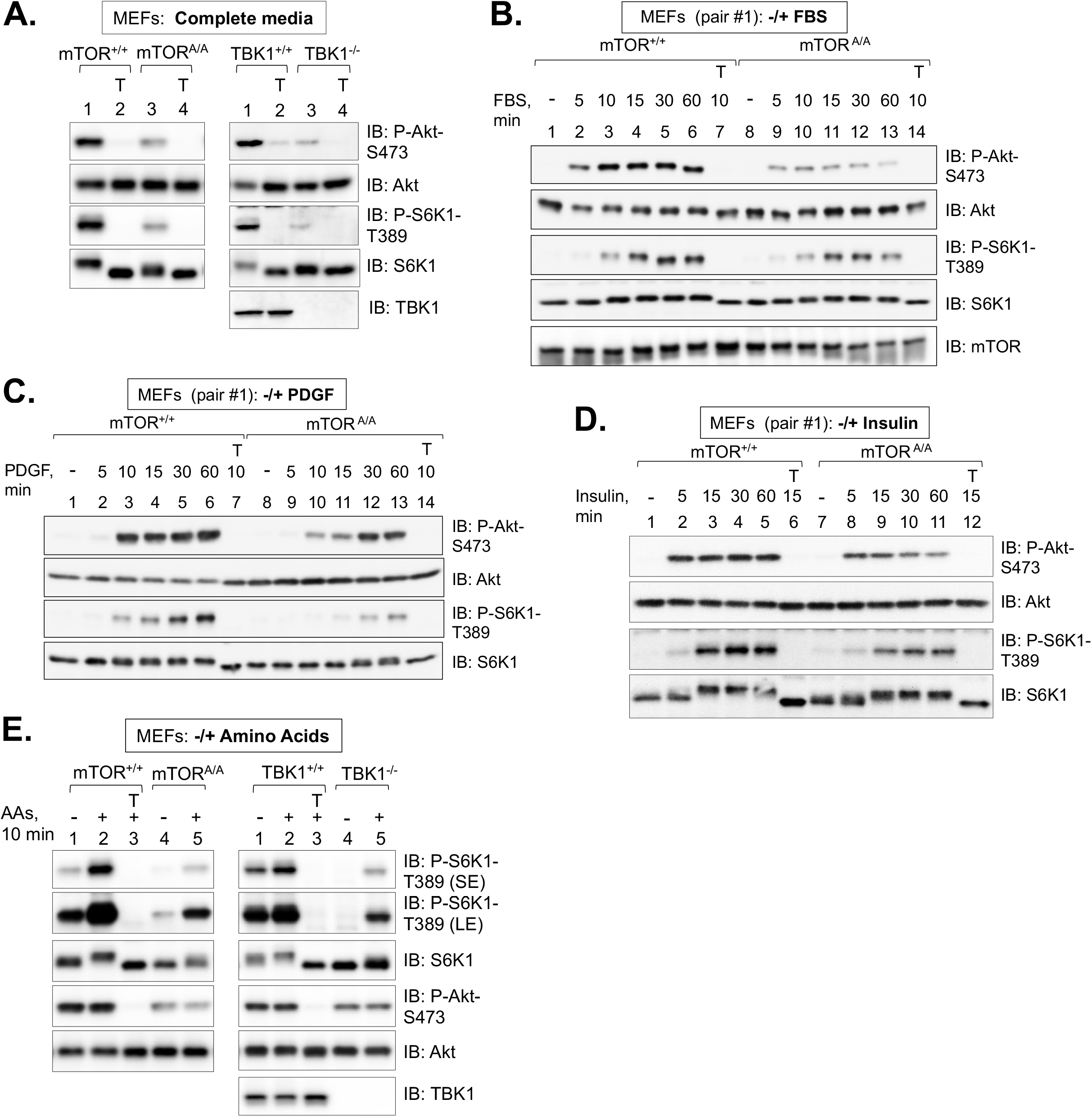
TBK1 and mTOR S2159 phosphorylation promote mTORC2 and mTORC1 signaling in response to diverse growth factors. **A. mTORC2 and mTORC1 signaling in mTOR S2159A knock-in MEFs and TBK1 null MEFs in complete media.** mTOR^+/+^ vs. mTOR^A/A^ MEFs (pair #1) and TBK1^+/+^ vs. TBK1^-/-^ MEFs were cultured in complete media (DMEM/FBS [10%]). At −80% confluency, cells were re-fed with complete media for 1.5 hr, treated without or with Torin1 (T) [100 nM] (30 min), and lysed. Whole cell lysates (WCLs) were immunoblotted with the indicated antibodies. **B. Effect of FBS on Akt P-S473 in mTOR S2159A knock-in MEFs.** mTOR^+/+^ and mTOR^A/A^ MEFs (pair #1) were serum starved overnight (20 hr), pre-treated with Torin1 (T) [100nM] (30 min.), and stimulated without (-) or with FBS [10% final] for the indicated times. WCLs were immunoblotted with the indicated antibodies. **C. Effect of PDGF on Akt P-S473 in mTOR S2159A knock-in MEFs.** mTOR^+/+^ and mTOR^A/A^ MEFs (pair #1) were serum starved overnight and treated as in 7B, except they were stimulated with PDGF [10 ng/mL]. WCLs were immunoblotted with the indicated antibodies. **D. Effect of insulin on Akt P-S473 in mTOR S2159A knock-in MEFs.** mTOR^+/+^ and mTOR^A/A^ MEFs (pair #1) were serum starved overnight and treated as in 7B, except they were stimulated with insulin [100 nM]. WCLs were immunoblotted with the indicated antibodies. **E. mTORC2 and mTORC1 signaling in mTOR S2159A knock-in MEFs and TBK1 null MEFs in the during amino acids deprivation and in response to acute amino acid stimulation.** mTOR^+/+^ vs. mTOR^A/A^ MEFs (pair #1) and TBK1^+/+^ vs. TBK1^-/-^ MEFs cultured in complete media (DMEM/FBS) were amino acid deprived in DMEM lacking all amino acids but containing 10% dialyzed FBS (dFBS) (50 min). MEFs were then stimulated with a mixture of 1x total amino acids (pH 7.4) (10 min.). WCLs were immunoblotted with the indicated antibodies. SE, short exposure; LE, long exposure.

As TBK1 knockout MEFs displayed reduced Akt S473 and S6K1 T389 phosphorylation upon amino acid stimulation following amino acid deprivation (14), we next examined a role for mTOR P-S2159 in control of these phosphorylation events in the absence or presence of amino acids. It is important to note that mTORC1 localization and activity change dynamically in response to amino acids levels, with the majority of studies indicating that mTORC2 does not function in amino acid sensing (30,61–66). We cultured mTOR^+/+^ vs. mTOR^A/A^ MEFs, as well as TBK1^+/+^ vs. TBKT^-/-^ MEFs, in DMEM lacking amino acids but supplemented with dialyzed FBS for 50 minutes; we then added back a 1x mixture of total amino acids (pH normalized to 7.4) for 10 minutes. Amino acids increased S6K1 P-T389 but not Akt P-S473 in wild type MEFs, as expected (Figure 7E) (DK 964; DK 961). mTOR^A/A^ and TBK1^-/-^ MEFs displayed reduced Akt P-S473 and S6K1 P-T389 in both the absence and presence of amino acids (Figure 7E). Similar to wild type MEFs, amino acids increased S6K1 P-T389 but not Akt P-S473 in the mutant MEFs. Taken together, these results indicate that amino acids activate mTORC1 but not mTORC2 signaling, and TBK1 and mTOR P-S2159 support both mTORC1 and 2 signaling in parallel to growth factor and amino acid signaling pathways.

### TBK1 activity and mTOR S2159 phosphorylation increase TLR3-mediated mTORC2 signaling in macrophages

Our prior work demonstrated that TBK1 and mTOR S2159 phosphorylation promote mTORC1 signaling in macrophages upon activation of TLR3 and 4, pathogen recognition receptors that activate TBK1 and IKKε (13). To investigate a role for TBK1 and mTOR P-S2159 in positive control of mTORC2 signaling in macrophages, we assayed how TBK1/IKKε inhibitors or *Mtor S2159A* knock-in modulated mTORC2 signaling to Akt in response to TLR3 activation with poly(I:C) (a viral dsRNA mimetic). In cultured RAW264.7 macrophages, poly(I:C) activated TBK1 (as monitored by increased TBK1 P-S172) and PI3K dependent mTORC2 signaling (as monitored by the sensitivity of Akt P-S473 to the class I PI3K inhibitor BYL-719 and the mTOR inhibitor Ku-0063794 (Figures 8A, S4) (IEG 69) (IEG 28). Inhibition of TBK1/IKKε with two different small molecules, amlexanox (Figures 8A, S4) (IEG 69) (IEG 28) or BX-795, (Figure S4)(IEG 28) also reduced poly(I:C) induced Akt P-S473, demonstrating that TBK1/IKKε activity positively controls mTORC2 signaling to Akt in RAW24.7 macrophages.

**Figure 8:**
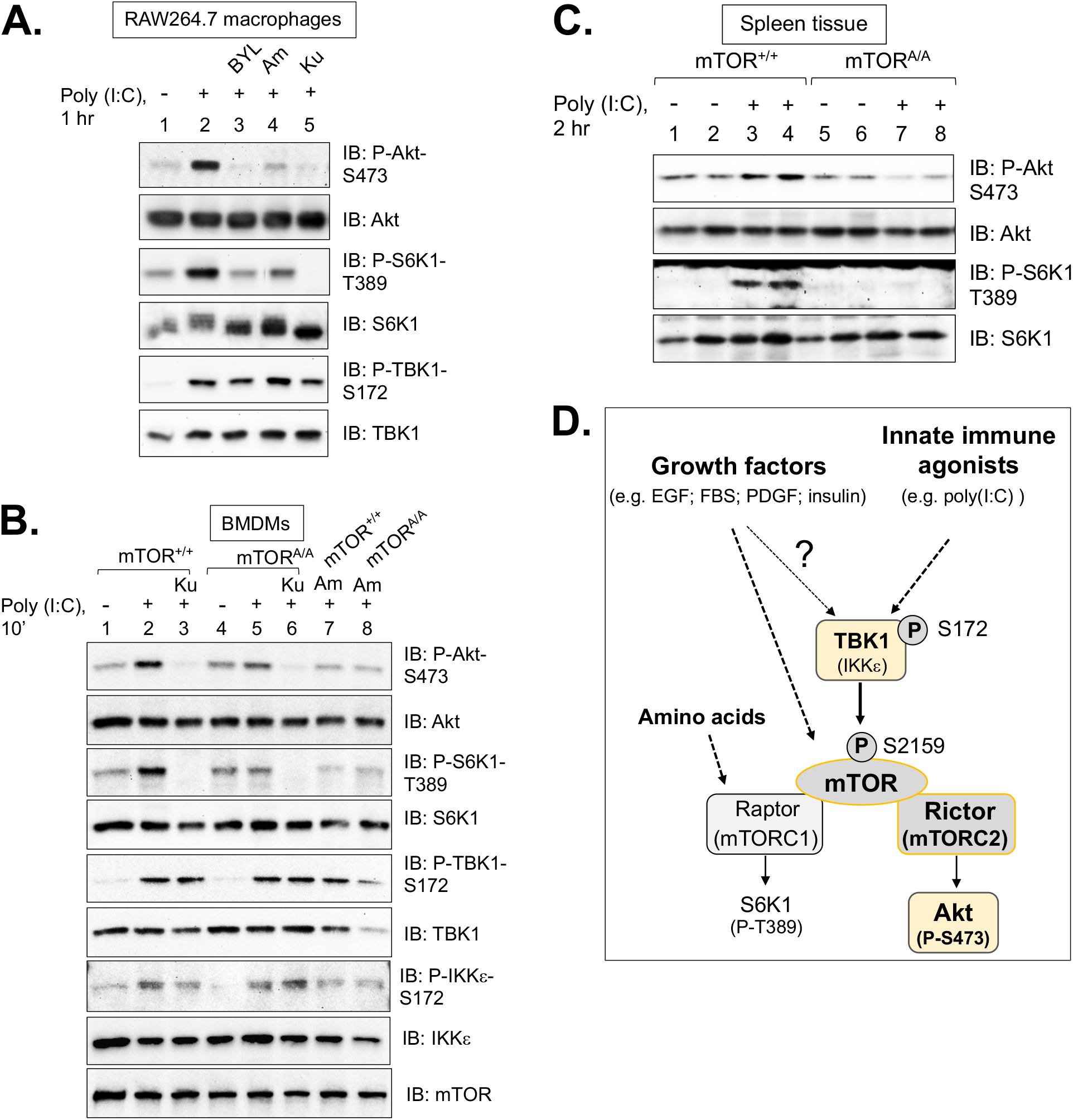
TBK1 activity and mTOR S2159 phosphorylation increase TLR3-mediated mTORC2 signaling in macrophages. **A. Effect of inhibiting PI3K (with BYL-719), TBK1 (with amlexanox), and mTOR (with Ku-0063794) on mTORC2 signaling in RAW264.7 macrophages in response to poly(I:C).** RAW264.7 macrophages cultured in complete media (DMEM/FBS) were pre-treated with BYL-719 [10 μM] (30 min), amlexanox [100 μM] (1 hr), or Ku-0063794 [100 nM] (30 min) and stimulated without (-) or with (+) poly(I:C) [30 μg/mL] (60 min). Whole cell lysates (WCLs) were immunoblotted with the indicated antibodies. **B. Effect of mTOR S2159A knock-in or amlexanox on mTORC2 signaling in primary BMDMs in response to poly(I:C).** Primary bone marrow derived macrophages (BMDMs) derived from *Mtor^+/+^* and *Mtor^A/A^* mice were cultured in complete media (DMEM/FBS), pre-treated with Ku-0063794 [100 nM] (30 min) or amlexanox [100 μM] (1 hr), and stimulated without (-) or with (+) poly(I:C) [30 μg/mL] (10 min). WCLs were immunoblotted with the indicated antibodies. **C. Effect of mTOR S2159A knock-in on mTORC2 signaling in mouse spleen tissue in response to poly(I:C) treatment *in vivo*.** *Mtor^+/+^* and *Mtor^A/A^* mice were fasted 5 hr and injected intraperitoneally with poly(I:C) [10 mg/kg-BW] (2 hr). Spleen tissue was isolated, homogenized, and analyzed by western blotting with the indicated antibodies. **D. Model.** See text for details.

We next isolated primary bone marrow-derived macrophages (BMDMs) from *Mtor^+/+^* and *Mtor^A/A^* mice. As expected, poly(I:C) increased TBK1 P-S172 and mTOR dependent Akt P-S473 in wild type BMDMs (Figure 8B) (CB 72). As in RAW264.7 macrophages, amlexanox suppressed Akt P-473 to a basal level (Figure 8B). Importantly, BMDMs from *Mtor^A/A^* mice displayed reduced poly(I:C) induced Akt P-S473 (Figure 8B), thus demonstrating a required role for mTOR S2159 phosphorylation in TLR3-mediated activation of mTORC2 signaling. Consistent with our prior work (13), amlexanox or *Mtor S2159A* knock-in reduced mTORC1 signaling in RAW264.7 macrophages and BMDMs (Figures 8A, 8B). These results demonstrate that TBK1 activity, mTOR activity, and mTOR S2159 phosphorylation are required for mTORC2 signaling to Akt upon activation of TLR3 in macrophages. Finally, to demonstrate a role for mTOR S2159 phosphorylation in activation of mTORC2 signaling by TLR3 *in vivo*, we injected *Mtor^+/+^* and *Mtor^A/A^* mice with poly(I:C) and harvested macrophage-rich spleen tissue. We found that spleen tissue from *Mtor^A/A^* mice displayed reduced Akt S473 phosphorylation in response to poly(I:C) (Figure 8C) (CB 233).

Taken together, these results demonstrate that TBK1 phosphorylates mTOR S2159 to activate mTORC2 directly and thus increase downstream signaling to Akt in cultured cells and *in vivo* (Figure 8D) (model cartoon). Moreover, they reveal that mTORC2 represents an essential link between TBK1 and Akt phosphorylation at physiological levels of protein expression. We find that in MEFs, basal TBK1 kinase activity signals in parallel to growth factors to augment mTORC2 (and mTORC1) activity, as EGF and other growth factors increased mTORC1/2 signaling in a TBK1 and mTOR P-S2159 dependent manner without increasing TBK1 S172 or mTOR S2159 phosphorylation. The relationship between growth factor signaling and TBK1 activity appears to be context dependent, however, as growth factors were shown recently to activate TBK1 (i.e., increase TBK1 P-S172) in lung cancer cells (15)(see Discussion). In macrophages, TLR3 signaling increases TBK1 and mTORC1/2 activity in a linear pathway (Figure 8D).

## Discussion

Several studies have demonstrated a positive role for TBK1 in Akt phosphorylation in various contexts. Increasing or decreasing TBK1 activity by various approaches in many cell types (e.g., MEFs, HEK293 cells, U2OS cells, HeLa cells, MNT1 melanoma cells, or HCT116 colorectal cancer cells) led to correspondingly similar changes in Akt S473 phosphorylation (14–17). These observations, together with evidence that mTORC2 serves as a major Akt S473 kinase (19–22) and our prior work that TBK1 directly activates mTORC1 (13), prompted us to investigate whether mTORC2 represents a missing link between TBK1 and Akt phosphorylation.

While it is challenging to demonstrate definitely that a kinase (e.g., TBK1) phosphorylates a substrate (e.g., mTOR) directly in intact cells rather than indirectly in a complex, several lines of evidence considered together support our model that TBK1 phosphorylates mTOR S2159 directly to increase mTORC2 activity and signaling to Akt (Figure 8D): TBK1 phosphorylates mTOR S2159 within mTORC2 *in vitro;* TBK1 and mTORC2 co-immunoprecipitate and TBK1^-/-^ MEFs display reduced mTOR P-S2159 in intact cells; TBK1^-/-^ MEFs and mTOR^A/A^ MEFS (lacking mTOR P-S2159) display reduced mTOR dependent Akt P-S473 and Akt P-T308 in response to EGF; Rictor^-/-^ MEFs stimulated with EGF bear extremely low Akt P-S473 that remains strongly Torin1 sensitive; TBK1 overexpression at low levels increases Akt P-S473 in a largely Torin1 sensitive manner; and finally, both TBK1^-/-^ MEFs and mTOR^A/A^ MEFs display reduced mTORC2 intrinsic catalytic activity in response to EGF, as measured by mTORC2 IVK assay and by Rictor-associated mTOR S2481 autophosphorylation. To support these results in immortalized cells, we also provide evidence that in primary macrophages in culture (i.e., BMDMs) and spleen tissue *in vivo*, mTOR S2159 phosphorylation is required for Akt S473 phosphorylation in response to TBK1 activation with poly(I:C). In addition, Zhao *et al*. found that adipocyte-specific knockout of TBK1 in diet-induced obese mice reduced Akt P-S473 in response to insulin in white adipose tissue (50). Taken together, these results argue that TBK1-mediated mTOR S2159 phosphorylation promotes mTORC2 signaling to Akt.

Several groups identified TBK1 as a direct Akt S473 and T308 kinase (16–18). We found that at physiological levels of TBK1 and Akt expression, the ability of growth factors or poly(I:C) to increase Akt S473 phosphorylation in a detectable manner required mTOR activity. mTOR inhibition also reduced Akt P-T308, a finding consistent with the observation that Akt S473 phosphorylation promotes and/or stabilizes Akt T308 phosphorylation (19,35,39). We found that overexpression of TBK1 increased Akt P-S473, similar to earlier work (16–18). This effect was largely dependent on mTOR activity at low to mid doses of TBK1 but displayed modest mTOR independence at higher doses. When both TBK1 and Akt were overexpressed, however, TBK1 increased Akt P-S473 in an mTOR independent manner. Taken together, these data indicate that at physiological levels of TBK1 and Akt expression, TBK1 increases Akt phosphorylation through mTORC2.

Pathological or unique physiological contexts may modify mechanisms governing Akt S473 and T308 phosphorylation. For example, tissue-specific knockout of Rictor or mTOR from cardiac or skeletal muscle failed to ablate Akt S473 phosphorylation (an unexpected finding) (17,67–69). Even more surprising, mTOR knockout cardiac muscle displayed elevated Akt P-S473 (17,69). This finding prompted Xie *et al*. to search for alternate Akt S473 kinases. Upon discovering TBK1 as an Akt S473 and T308 kinase, they noted that mTOR knockout cardiac muscle displayed elevated TBK1 expression (17). Thus, in this context of elevated TBK1 expression, TBK1 may phosphorylate Akt directly. Pathological contexts may also render Akt S473 phosphorylation TBK1 independent. For example, TBK1 inactivation reduced S6K1 T389 but not Akt S473 phosphorylation in response to growth factors in A549 human lung adenocarcinoma cells or primary mouse lung cancer epithelial cells (15). Therefore, we propose that in stress-related contexts (e.g., knockout of the essential kinase mTOR), oncogenic contexts, or other pathological contexts, cells and tissues may re-wire signaling and metabolism by upregulating expression of TBK1/ IKKε (or other Akt S473 kinases). Such an adaptive response may in turn enable TBK/ IKKε (or other kinases) to directly phosphorylate Akt S473 and/or T308, thus increasing Akt activity in order to mitigate the pathologic insult and improve metabolic homeostasis and/or provide a proliferative or survival advantage.

Mechanisms governing regulation of TBK1 kinase activity in non-immune cells remain poorly defined. Our results indicate that in MEFs, growth factors increase mTORC2 signaling in a TBK1 and mTOR P-S2159 dependent manner without increasing TBK1 S172 phosphorylation (i.e., TBK1 activity) or mTOR S2159 phosphorylation. We therefore conclude that in this context, the basal kinase activity of TBK1 promotes mTORC2 signaling in parallel to growth factors. It is important to note, however, that stimulation of human A549 lung cancer cells and primary mouse lung cancer cells with several different growth factors (e.g., EGF; FBS; insulin) increased TBK1 S172 phosphorylation, which required TBKBP1 (TBK1 binding protein-1), a TBK1 adaptor protein (15). We speculate that differential expression of numerous TBK1 adaptors explains differences in cellular context that underlies regulation of TBK1 activity by growth factors. These adaptors may also differentially control TBK1 subcellular localization and/or TBK1 substrate preference (7,46,70,71).

Taken together, our results identify TBK1 as a direct activator of mTORC2 in physiological contexts, which expands our limited understanding of mTORC2 regulation. In addition, they establish the TBK1-mTORC2 pathway as a potential target for therapeutic intervention to treat cancer and obesity linked metabolic disorders.

## Experimental Procedures

### Materials

General chemicals were from Thermo Fisher or Sigma. Protein A- and G-Sepharose Fast Flow were from GE Healthcare; NP40, Brij35, and CHAPS (3-[(3-cholamidopropyl)-dimethylammonio]-1-propanesulfonate) detergents were from Pierce; cOmplete Protease Inhibitor Cocktail (EDTA-free) tablets were from Millipore Sigma (#11836170001); Immobilon-P polyvinylidene difluoride (PVDF) membrane (0.45 μM) was from Millipore; Bradford Reagent for protein assays was from Bio Rad (#5000001); reagents for enhanced chemiluminescence (ECL) were from either Alkali Scientific (Bright Star) or Advansta (Western Bright Sirius HRP substrate). Recombinant active GST-TBK1 was from Thermo Fisher/ Life Technologies (#PV3504); recombinant His-Akt1 was from EMD Millipore (#14-279); recombinant GST-mTORf (containing a 30 fragment of mTOR encoding amino acids 2144-2175) was generated as described (*13*).

### Antibodies

The following antibodies from Cell Signaling Technology (CST) were used in this study (all rabbit polyclonal antibodies, unless otherwise noted): Akt (#9272); Akt P-S473 (#4060; rabbit mAb D9E XP); Akt P-T308 (#4056, rabbit mAb 244F9); TBK1 (#3013; or #3504, rabbit mAb D1B4); TBK1 P-S172 (#5483, rabbit mAb D52C2 XP); S6K1 (#9202); S6K1 P-T389 (#9205; #9234, rabbit mAb 108D2; #9206, mouse mAb IA5); mTOR (#2972); MAPK (#9102); MAPK P-T202/Y204 (#4370, rabbit mAb D13.14.4E XP); GST (#2624; mouse mAb 26H1); IgG-conjugated Sepharose beads (#3423); Rictor-conjugated Sepharose beads (#5379). The following antibodies were from other commercial vendors: mTOR P-S2481 (MilliporeSigma #09-343); Myc (Millipore Sigma #05-419, mouse mAb 9E10); HA.11 (BioLegend # 901513, mouse mAb 1612B). The following custom, anti-peptide polyclonal antibodies were generated by us in-house with assistance from a commercial vendor: Rictor (amino acids 6-20; Covance; as described (*72,73*)); mTOR (amino acids 221-237, Covance; as described (*72,73*)); mTOR P-S2159 (Millipore; as described (*13*)).

### Plasmids

pcDNA3/Flag-TBK1 wild type (WT) and pcDNA3/Flag-TBK1kinase dead (K38M) were obtained from Dr. A. Saltiel (UCSD School of Medicine, Institute for Diabetes and Metabolic Health; San Diego, CA). pcDNA3/Flag-HA-Akt was obtained from Addgene (#9021). pRK5/Myc-mTOR WT was originally from Dr. D. Sabatini (MIT and the Whitehead Institute, Boston, MA) and obtained via Addgene (#1861). pRK5/Myc-mTOR S2159A was generated via site-directed mutagenesis as described (*13*)). pCI/HA-Rictor was from Dr. E. Jacinto (Rutgers University, New Brunswick, NJ).

### Cell culture, transfection, and drug treatments

All cell lines used in this study (i.e., MEFs, HEK293, RAW264.7 murine macrophages, and primary mouse BMDMs) were cultured in DMEM that contained high glucose [4.5 g/L], glutamine [584 mg/L], and sodium pyruvate [110 mg/L] (Thermo Fisher/ Life Technologies) supplemented with 10% fetal bovine serum (FBS) (Gibco/Invitrogen). Note that dialyzed FBS was used to culture the RAW264.7 macrophages and BMDMs. Cells were incubated at 37°C in a humidified atmosphere containing 5% CO_2_. HEK293 cells (from ATCC) were transfected using Mirus Trans-It LT1 in accordance with manufacturer’s instructions and lysed ~24 to 48 hr. post-transfection. To stimulate cells with growth factors, MEFs and HEK293 cells were first serum starved via incubation in DMEM containing 20 mM Hepes pH 7.2 overnight, ~20 hr. The cells were then stimulated with EGF [25 or 50 ng/mL] (Sigma Aldrich #E9644 and #E4127), FBS [10% final], PDGF [10 ng/mL] (EMD Millipore #GF149), or insulin [100 nM] (Thermo Fisher/ Life Technologies #12585) for various amounts of time (0-60 min.). To stimulate macrophages with innate immune agonists, RAW264.7 macrophages and BMDMs cultured in complete DMEM (i.e., DMEM containing dialyzed FBS [10%]) were treated with poly(I:C) [30 ug/ mL] (Sigma Aldrich #P1530) or ultrapure LPS [100 ng/ml] (InVivo Gen #tlrl-3pelps). Cells were treated with the following drugs: Torin1 [100 nM] (30 min) (shared by Dr. D. Sabatini), Ku-0063794 [100 nM] (30 min) (Tocris #3725); amlexanox [100 uM] (1-2 hr) (EMD Millipore #SML0517); BX-795 [10 uM] (30 min)(Millipore CalBiochem #204011)(30 min); BYL-719 [10 uM] (30 min) (Selleck; #1020). To amino acid deprive and stimulate cells, MEFs were first incubated for 50 min in RPMI media lacking all amino acids (Millipore Sigma; # R8758) but supplemented with dialyzed FBS [10%]. The cells were then acutely stimulated with amino acids for 10 min by adding a mixture of total amino acids (RPMI 1640 Amino Acid Solution) (Millipore Sigma; # R7131) to a final concentration of [1x] (the concentration of amino acids in RPMI medium). Note that the pH of this amino acid solution is quite basic, and thus the pH of this solution was first normalized to 7.4 prior to addition to cells in order to maintain physiological pH. As the RPMI 1640 Amino Acid Solution lacks glutamine, the amino acid mixture was supplemented with L-glutamine (Millipore Sigma; #59202C) prior to pH normalization and addition to cells.

### Cell lysis, immunoprecipitation, and immunoblotting

Cells were washed twice with ice-cold PBS pH 7.4 and scraped into ice-cold lysis buffer A (10 mM KPO4 pH 7.2; 1 mM EDTA; 5 mM EGTA; 10 mM MgCl_2_; 50 mM β-glycerophosphate; 1 mM sodium orthovanadate [Na_3_VO_4_]; a cocktail of protease inhibitors) containing NP-40 [0.5%] and Brij35 [0.1%], as described (*73*). To maintain detergent sensitive mTOR-partner protein interactions during Rictor or mTOR immunoprecipitation, cells were lysed in ice-cold buffer A containing mild CHAPS [0.3%] detergent. Lysates were spun at 13,200 rpm for 5 min at 4°C, and the post-nuclear supernatants were collected and incubated on ice (15 min). Bradford assay was used to normalize protein levels for immunoblot or immunoprecipitation analysis. For immunoprecipitation, whole cell lysates were incubated with antibodies for 2 hr. at 4°C, followed by incubation with Protein G- or A-Sepharose beads for 1 hr. Sepharose beads were washed three times in lysis buffer and resuspended in 1x sample buffer. Samples were resolved on SDS-PAGE and transferred to PVDF membranes in Towbin transfer buffer containing 0.02% SDS, as described (*73*). Immunoblotting was performed by blocking PVDF membranes in Tris-buffered saline (TBS) pH 7.5 with 0.1% Tween-20 (TBST) containing 3% non-fat dry milk, as described (*73*), and incubating the membranes in TBST with 2% bovine serum albumin (BSA) containing primary antibodies or secondary HRP-conjugated antibodies. Blots were developed by ECL and detected digitally with a Chemi-Doc-It System (UVP).

### Lentiviral transduction

TBK1^-/-^ MEFs stably expressing Flag-TBK1 (wild type or kinase dead K38M) were generated by lentiviral transduction. Flag-TBK1 was subcloned into a modified lentiviral vector, pHAGE-Puro-MCS (pPPM) (74) (modified by Amy Hudson (Medical College of Wisconsin) to include an expanded multiple cloning site, MCS). Lentivirus particles were packaged in HEK293T cells by co-transfection with empty pPPM vector or pPPM/Flag-TBK1 together with pRC/Tat, pRC/Rev, pRC/gag-pol and pMD/VSV-G using Mirus TransIT-LT1 transfection reagent. Supernatants containing viral particles were collected 48 hr. post transfection and filtered through a 0.45 μm filter. TBK1^-/-^ MEFs were infected with fresh viral supernatants containing 8 μg/ml polybrene. 24 hr. post infection, cells were re-fed with DMEM/10% FBS supplemented with 10 μg/ml puromycin for 2-3 days to select for stably transduced cells, trypsinized, and plated at limiting dilution in order to isolate clones originating from single cells. TBK1^-/-^ MEF clones transduced with wild type Flag-TBK1 lentivirus were analyzed for expression of exogenous Flag-TBK1 relative to expression of endogenous TBK1 found in wild type MEFs. TBK1^-/-^ MEF clones expressing Flag-TBK1 at a level similar to endogenous TBK1 were chosen for analysis. Alternately, TBK1^-/-^ MEFs transduced with wild type or kinase dead (K38M) Flag-TBK1 were selected in puromycin and pooled for analysis. Rictor^-/-^ MEFs (from Dr. E. Jacinto; Rutgers University, New Brunswick, NJ) stably expressing vector control or HA-Rictor were generated by lentiviral transduction and stable selection, as described (35). These rescued lines were maintained in DMEM/FBS containing puromycin [8 μg/mL]. To knockdown TBK1, Rictor^-/-^ MEFs were infected with lentiviral particles encoding an shRNA targeting TBK1 (Sigma) (mouse TBK1 # TRCN0000323444; non-targeting # SHC016V) and then selected in puromycin [8 μg/mL] for 4 days.

### In vitro kinase assays

#### Phosphorylation of recombinant His-Akt1 or GST-mTORf by recombinant active TBK1

*In vitro* kinase (IVK) assays were performed by incubating recombinant His-Akt1 [50 ng] or GST-mTOR_f_ [50 ng] substrates with recombinant active GST-TBK1 [50 ng] in kinase buffer containing 25 mM Tris-HCl pH 7.5, 10 mM MgCl_2_, 1 mM DTT, and 200 μM ATP. Reactions were incubated at 30°C for 30 min and stopped by addition of sample buffer followed by incubation at 95°C for 5 min. Samples were resolved on SDS-PAGE, transferred to PVDF membrane, and immunoblotted. The phosphorylation of His-Akt1 was measured using anti-Akt P-S473 antibodies, and the phosphorylation of GST-mTORf was monitored using anti-mTOR P-S2159 antibodies. For certain IVK reactions, recombinant kinases were pre-incubated with BX-795 [10 μM] in kinase buffer on ice for 30 min.

#### Phosphorylation of Myc-mTOR or Rictor-associated mTOR isolated from cells by recombinant active TBK1

HEK293 cells were transfected with Myc-mTOR wild type or mutant S2159A, and lysates were immunoprecipitated with anti-Myc antibodies. Myc-mTOR S2159A was generated by QuikChange mutagenesis, as described previously (12). Alternately, Rictor from non-transfected HEK293 cells was immunoprecipitated with anti-Rictor antibodies. After washing in lysis buffer and kinase buffer, the immune complexes (containing substrate) were incubated with recombinant active GST-TBK1 [50 ng] in kinase buffer and ATP, as described above. The phosphorylation of Myc-mTOR or Rictor-associated mTOR was monitored using anti-mTOR P-S2159 antibodies.

#### Phosphorylation of His-Akt1 by cellular mTORC2 (i.e., mTORC2 IVKs)

mTORC2 *in vitro* kinase (IVK) assays was performed as described (35,75). Briefly, Rictor was immunoprecipitated from serum-starved MEFs pre-treated without or with Torin1 [100 nM] (30 min) and then stimulated without or with EGF [50 ng/mL] (10 min) (~one 10 cm plate for each immunoprecipitate). After washing in lysis buffer and kinase buffer, the immune complexes (containing mTOR kinase) were incubated with ATP [250 μM] and recombinant His-Akt1 [100 ng/reaction] in kinase buffer containing 25 mM HEPES, 100 mM potassium acetate, and 1 mM MgCl_2_ at 30°C for 30 min. Certain reactions were also pre-treated with Torin1 [100 nM10 μM] in kinase buffer on ice for 30 min prior to initiating the IVK reaction. Immunoprecipitates were pre-incubated with Torin1 [100 nM] on ice (30 min) and then were incubated with ATP [250 μM] and recombinant His-Akt1 [100 ng/reaction] in kinase buffer containing 25 mM HEPES, 100 mM potassium acetate, and 1 mM MgCl_2_, at 30°C for 30 min.

### Co-immunoprecipitation of TBK1 and mTORC2

TBK1^+/+^ or TBK1^-/-^ MEFs were lysed in buffer A containing CHAPS [0.3%] detergent. Protein levels in each cell type were normalized after performing protein assays with the Bradford assay. IgG control Sepharose beads (CST) or Rictor-conjugated Sepharose beads (CST) were washed 2x with PBS, blocked ~1 hr in PBS containing 2% bovine serum albumin (BSA) to reduce non-specific binding, and then washed in PBS 2x more. Whole cell lysates were then added to the washed beads, rotated at 4° overnight, washed 3x in lysis buffer, and resuspended in 1x SDS-PAGE sample buffer.

### Mtor S2159A knock-in mice

mTOR knock-in S2159A mice (*Mtor^A/A^*) were generated using CRISPR-Cas9 genome editing technology and genotyped, as described (13). Mice were housed in a specific pathogen-free facility with a 12-hour light/ 12-hour dark cycle and given free access to food and water. All animal use was in compliance with Institutional Animal Care and Use Committee (IACUC) at the University of Michigan.

### Isolation and immortalization of MEFs

Male and female heterozygous *Mtor^+/A^* mice were mated, and MEFs from plugged females were isolated on day 13.5 of pregnancy, generally as described (76,77). Dissected embryos were washed with 3x in PBS pH 7.4 and minced with fine scissors in the presence of trypsin-EDTA. The minced tissue was triturated with 5 mL serological and fine-tip Pasteur pipettes to prepare a homogenate and taken into a 15 mL conical tube containing 8 mL DMEM. The homogenate was centrifuged 4 min at 300g, and the supernatant was discarded. The pellet was washed 1x in PBS pH 7.4 and resuspended in complete medium (DMEM/ FBS [10%] containing 50 U/mL penicillin and 50 μg/mL streptomycin). The resuspended cells were transferred to a 10 cm culture dish with fresh complete medium and incubated at 37°C in a humidified atmosphere containing 5% CO_2_. The MEFs were washed 1x with PBS pH 7.4, detached with 0.05% trypsin-EDTA, centrifuged 4 min at 300g, and transferred to a 15 cm culture dish. At confluency, the MEFs were trypsinized and aliquoted to multiple cryovials for longterm storage in liquid N_2_. MEFs were immortalized spontaneously through serial passaging. As they reached confluency in 15 cm culture dishes following isolation, primary MEFs were plated into 10 cm culture dishes at a 1:9 split ratio into 10 ml complete medium (passage #2). Upon reaching confluency, the MEFs were again passaged into 10 cm culture dishes with a split ratio of 1:3. This process was repeated every 3-4 days until reaching senescence (~passage 5-7). Culture media was replaced every 3-4 days, and the cells were trypsinized and transferred into a new culture dish once a week. Once the MEFs began proliferating, they were passaged with a 1:3 split ratio every 3-4 days until cell number doubled every ~24-30 hrs. MEFs were genotyped as described (13) using dissected head tissue.

### Isolation of primary bone marrow derived macrophages (BMDMs)

Bone marrow from 8-14 wk old *Mtor^+/+^* and *Mtor^A/A^* mice was harvested by flushing femora and tibiae with ice-cold PBS pH 7.4 using a 30G needle under sterile conditions. Bone marrow cells were suspended in MEM with L-glutamine supplemented with 10% HI-FBS, 50 U/ml penicillin, 50 μg/ml streptomycin, and 20 ng/ml M-CSF (R&D Systems; #416-ML) and plated into 6-well tissue culture plates. Cells were incubated at 37°C in 5% CO_2_, and medium was replaced every other day until day 5, at which point the monocytes had differentiated into macrophages. Macrophages were studied at ~80% confluency.

### In vivo poly(I:C) treatment of mice

*Mtor^+/+^* and *Mtor^A/A^* mice (mostly C57BL6) (12 wks old) fed a normal chow diet were fasted (5 hr) and injected with saline or poly (I:C) [10 mg/kg-BW] (2 hr). Spleen tissue was isolated, homogenized, and analyzed by western blotting. A motorized tissue homogenizer (Tissue Ruptor, Qiagen) was used to homogenize whole spleen in 1 ml ice-cold RIPA buffer that contained protease and phosphatase inhibitors. Lysate was incubated on ice (10 min), centrifuged at 13,200 rpm (15 min, 4°C), and supernatant was collected. DC (Detergent Compatible) protein assay (Bio-Rad; #5000111) was used to standardize protein amount for immunoblot analysis.

### Western blot editing, quantification, and statistical analysis

Western blot images were prepared for publication using Adobe Photoshop. Only the parameters levels, brightness, and contrast were employed to fine-tune exposure time and band sharpness. Importantly, these parameters were adjusted equivalently across the entire blot, and the final image shown reflects the raw image. In certain panels, thin dotted lines indicate excision of an irrelevant lane(s) from a western blot. Western blot signals were quantitated using FIJI to measure integrated densities of protein bands. The ratios of phosphorylated proteins over cognate total protein were calculated and normalized as indicated in the figure legends. Statistical significance was tested using paired Student’s t-test assuming equal variances. Error bars represent either standard deviation (SD) or standard error of the mean (SEM), as indicated in the figure legend.

## Abbreviations used frequently in this study

TBK1: (TANK-binding kinase 1);
IKKε: Iκβ kinase ε;
mTOR: (mechanistic target of rapamycin);
mTORC: (mTOR complex);
mTORC2: (mTOR complex 2);
mTORC1: (mTOR complex 1);
Rictor: (rapamycin-insensitive companion of mTOR);
Raptor: (regulatory-associated protein of mTOR);
PI3K: phosphatidylinositol 3’ kinase;
IRF3/7: interferon regulatory factor 3/7;
IFNβ: interferon β

## *Supplementary information

This article contains supporting figures, S1-S4.

## Acknowledgments

We thank Dr. Alan R. Saltiel (UCSD School of Medicine, Institute for Diabetes and Metabolic Health; San Diego, CA) for sharing Flag-TBK1 plasmids (wild type and kinase dead) and RAW264.7 macrophages, as well as sharing information regarding TBK1 biology. We thank Dr. E. Jacinto (Rutgers University, New Brunswick, NJ) for sharing the plasmid pCI/HA-Rictor and Rictor^-/-^ MEFs.

We thank Dr. D. Sabatini (Whitehead Institute, MIT, Boston, MA) for sharing Torin1. We thank Dr. Ben Allen (University of Michigan Medical School, Dept. of Cell and Developmental Biology, Ann Arbor, MI) for assistance with isolation of mouse embryonic fibroblasts (MEFs). We also thank Dr. Daniel Lucas (Cincinnati Children’s Hospital) for assistance with isolation of bone marrow-derived macrophages (BMDMs).

## Funding

This work was supported by grants to DCF from the NIH (R01-DK-103877) and from the American Diabetes Association (ADA) (Basic Science Grant #1-12-BS-49). The work was also supported by the Molecular Genetics Core of the Michigan Diabetes Research Center (MDRC) (#P30-DK020572; NIH-NIDDK).

## Supplementary Figures

**Figure S1 (related to Figure 1).**
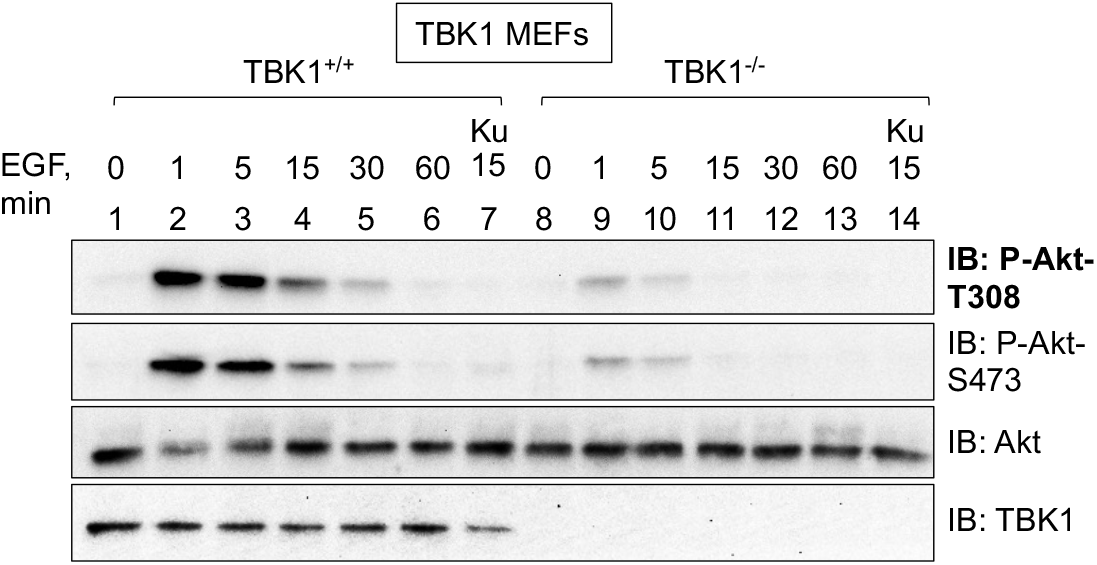
EGF stimulated Akt T308 phosphorylation in TBK1 null MEFs. TBK1^+/+^ and TBK1^-/-^ MEFs were serum starved overnight (20 hr), pre-treated with Torin1 (T) [100nM] (30 min), and stimulated without (-) or with (+) EGF [50 ng/mL] for the indicated times. Whole cell lysates were immunoblotted with the indicated antibodies.

**Figure S2 (related to Figure 4).**
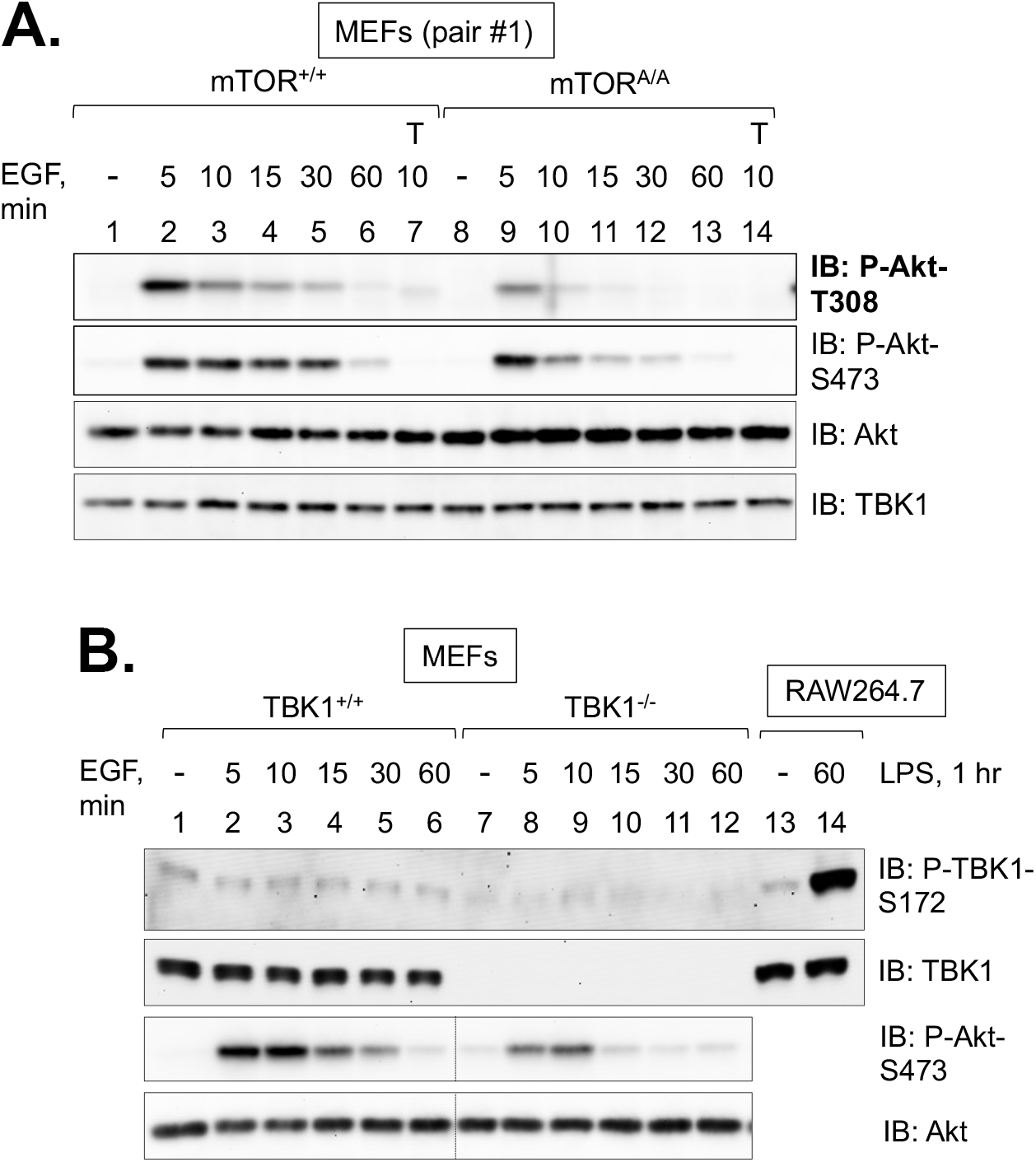
**S2A. Effect of EGF on Akt T308 phosphorylation in wild type and mTOR S2159A knock-in MEFs.** Immortalized mTOR^+/+^ and mTOR^A/A^ MEFs were serum starved overnight (20 hr), pre-treated with Torin1 (T) [100nM] (30 min), and stimulated without (-) or with (+) EGF [50 ng/mL] for the times indicated. Whole cell lysates (WCLs) were immunoblotted with the indicated antibodies. **S2B. Effect of EGF on TBK1 S172 phosphorylation in TBK1 MEFs.** TBK1^+/+^ and TBK1^-/-^ MEFs were serum starved overnight and stimulated with EGF for the times indicated, as in S2A. RAW264.7 macrophages in complete media were stimulated without (-) or with LPS [100 ng/mL] (1 hr) to serve as a positive control for TBK1 P-S172 western blotting. WCLs from MEFs and RAW264.7 macrophages were resolved on the same gel and immunoblotted with the indicated antibodies with protein amounts normalized between the two cell types.

**Figure S3 (related to Figure 7).**
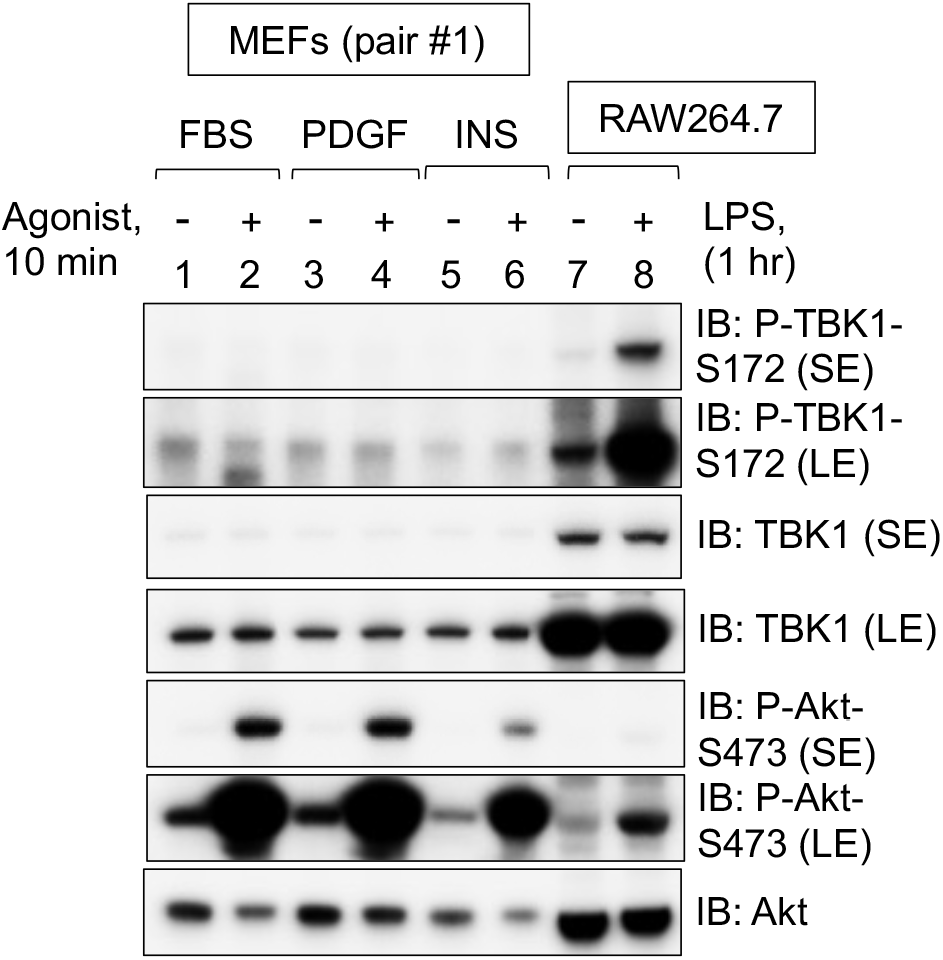
Effect of diverse growth factors on TBK1 P-S172 in mTOR MEFs. mTOR^+/+^ and mTOR^A/A^ MEFs (pair #1) were serum starved overnight and stimulated without (-) or with (+) FBS [10%], PDGF [10 ng/mL], or insulin [100 nM] (10 min). RAW264.7 macrophages in complete media were stimulated without (-) or with (+) LPS [100 ng/mL] (1 hr) to serve as a positive control for TBK1 P-S172 western blotting. Whole cell lysates from MEFs and RAW264.7 macrophages were resolved on the same gel and immunoblotted with the indicated antibodies. Note that protein amounts were not normalized between the two cell types.

**Figure S4 (related to Figure 8).**
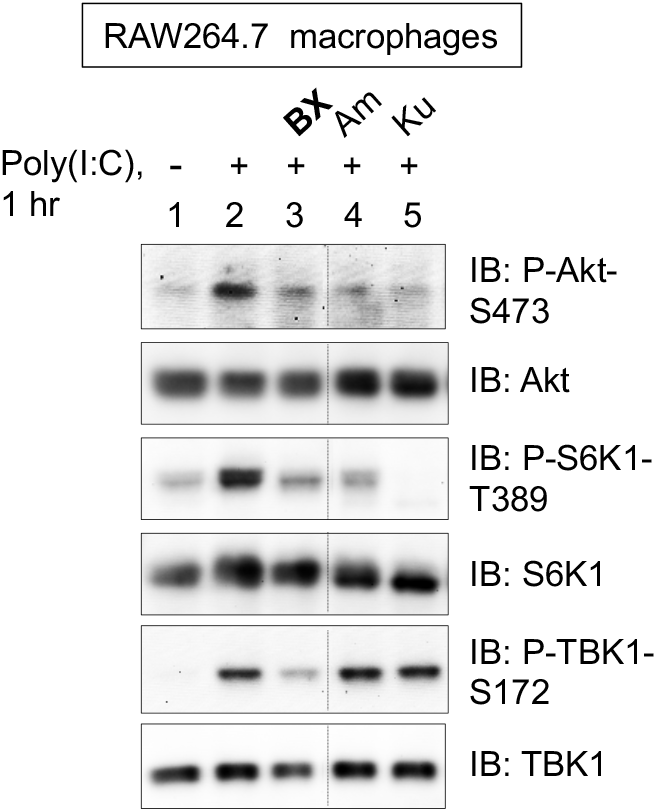
Effect of the TBK1/IKKε inhibitor BX-795 on mTORC2 signaling in RAW264.7 macrophages in response to poly(I:C). RAW264.7 macrophages cultured in complete media (DMEM/FBS) were pretreated with the TBK1/IKKε inhibitor BX-795 (BX)[10 μM] (30 min) as well as amlexanox (Am) [100 μM] (1 hr) and Ku-0063794 (Ku) [100 nM] (30 min) and stimulated without (-) or with (+) poly(I:C) [30 μg/mL] (60 min). Whole cell lysates were immunoblotted with the indicated antibodies.

## Notes

The authors declare that they have no conflicts of interest with the contents of this article.

